# Changes in intra- and interlimb reflexes from forelimb cutaneous afferents after staggered thoracic lateral hemisections during locomotion in cats

**DOI:** 10.1101/2024.04.23.590723

**Authors:** Stephen Mari, Charly G. Lecomte, Angèle N. Merlet, Johannie Audet, Sirine Yassine, Rasha Al Arab, Jonathan Harnie, Ilya A. Rybak, Boris I. Prilutsky, Alain Frigon

**Author notes:** Corresponding author: Alain Frigon, PhD.

## Abstract

In quadrupeds, such as cats, cutaneous afferents from the forepaw dorsum signal external perturbations and send signals to spinal circuits to coordinate the activity in muscles of all four limbs. How these cutaneous reflex pathways from forelimb afferents are reorganized after an incomplete spinal cord injury is not clear. Using a staggered thoracic lateral hemisections paradigm, we investigated changes in intralimb and interlimb reflex pathways by electrically stimulating the left and right superficial radial nerves in seven adult cats and recording reflex responses in five forelimb and ten hindlimb muscles. After the first (right T5-T6) and second (left T10-T11) hemisections, forelimb-hindlimb coordination was altered and weakened. After the second hemisection, cats required balance assistance to perform quadrupedal locomotion. Short-, mid- and long- latency homonymous and crossed reflex responses in forelimb muscles and their phase modulation remained largely unaffected after staggered hemisections. The occurrence of homolateral and diagonal mid- and long-latency responses in hindlimb muscles evoked with left and right superficial radial nerve stimulation was significantly reduced at the first time point after the first hemisection, but partially recovered at the second time point with left superficial radial nerve stimulation. These responses were lost or reduced after the second hemisection. When present, all reflex responses, including homolateral and diagonal, maintained their phase-dependent modulation. Therefore, our results show a considerable loss in cutaneous reflex transmission from cervical to lumbar levels after incomplete spinal cord injury, albeit with preservation of phase modulation, likely affecting functional responses to external perturbations.

**Key points:**

- Cutaneous afferent inputs coordinate muscle activity in the four limbs during locomotion when the forepaw dorsum contacts an obstacle.
- Thoracic spinal cord injury disrupts communication between spinal locomotor centers located at cervical and lumbar levels, impairing balance and limb coordination.
- We investigated cutaneous reflexes from forelimb afferents during quadrupedal locomotion by electrically stimulating the superficial radial nerve bilaterally, before and after staggered lateral thoracic hemisections in cats.
- We showed a loss/reduction of mid- and long-latency homolateral and diagonal reflex responses in hindlimb muscles early after the first hemisection that partially recovered with left superficial radial nerve stimulation, before being reduced after the second hemisection.
- Targeting cutaneous reflex pathways from forelimb afferents projecting to the four limbs could help develop therapeutic approaches aimed at restoring transmission in ascending and descending spinal pathways.

**Figure Abstract:** 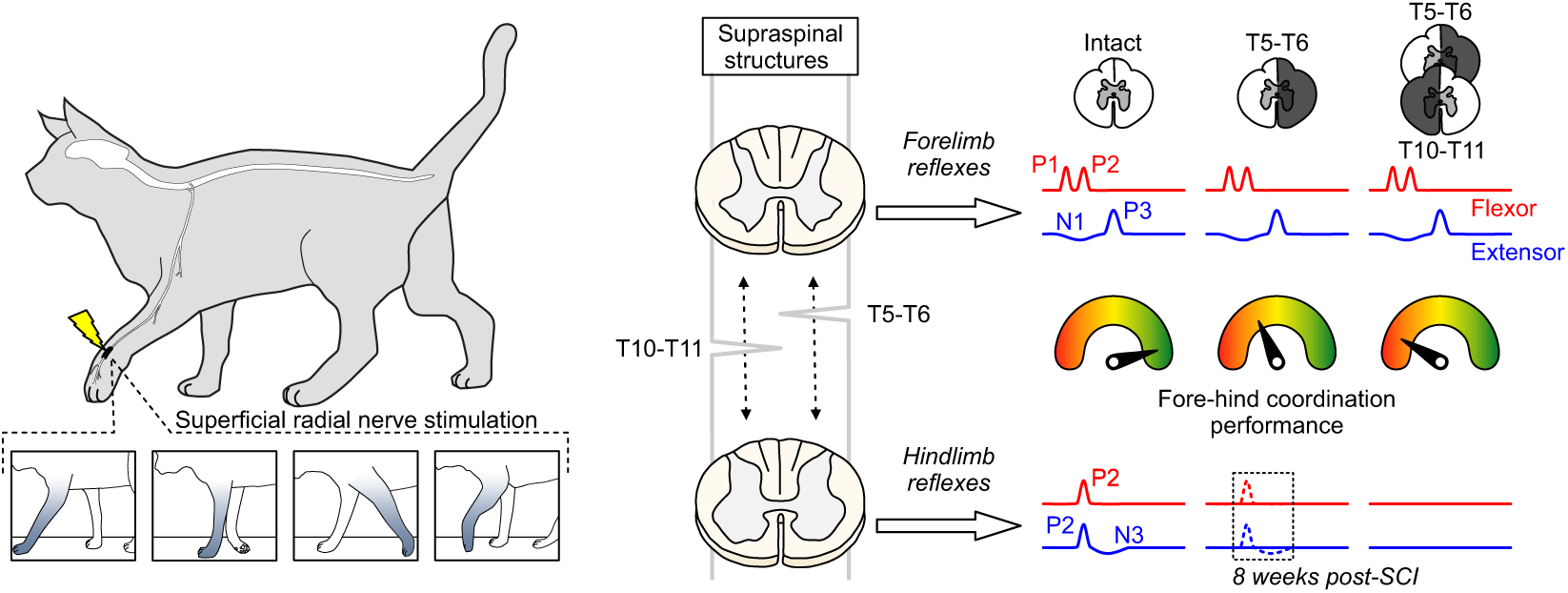

Contacting an obstacle during locomotion activates cutaneous afferents to maintain balance and coordinate all four limbs. Spinal cord injuries disrupt neural communications between spinal networks controlling the fore- and hindlimbs, impairing balance and limb coordination. Cutaneous reflex pathways can be used to develop therapeutic approaches for restoring ascending and descending transmission to facilitate locomotor recovery.

## INTRODUCTION

Locomotion, inputs from the skin provide information on the external environment, such as characteristics of the terrain and obstacles encountered (Rossignol *et al*., 2006; Pearcey & Zehr, 2019*a*; Frigon *et al*., 2021). For instance, in cats and humans, cutaneous afferents play an important role in modifying limb trajectory and maintaining balance during locomotion when the foot/hindpaw dorsum, innervated by the superficial peroneal (**SP**) nerve, contacts an obstacle during the swing phase, termed the stumbling corrective reaction (Forssberg *et al*., 1977; Prochazka *et al*., 1978; Forssberg, 1979; Duysens & Loeb, 1980; Wand *et al*., 1980; Schillings *et al*., 1996; Van Wezel *et al*., 1997; Zehr *et al*., 1997; Quevedo *et al*., 2005*b*, 2005*a*). In quadrupeds, such as cats, stimulating the dorsum of the forepaw or the superficial radial (**SR**) nerve also elicits a stumbling corrective reaction that alters forelimb trajectory to move it away and over a simulated obstacle (Miller *et al*., 1977; Matsukawa *et al*., 1982; Drew & Rossignol, 1985, 1987; Shimamura *et al*., 1990; Fuwa *et al*., 1991; Hurteau *et al*., 2018; Mari *et al*., 2023). In cats and humans, electrically stimulating the SP and SR nerves evoke short-, mid- and long-latency inhibitory and/or excitatory cutaneous reflex responses in muscles of the four limbs that are modulated with task and phase during locomotion (Haridas & Zehr, 2003; Mari *et al*., 2023). Mid- and long-latency responses are thought to involve supraspinal contributions, but also polysynaptic spinal pathways (Fuwa *et al*., 1991; LaBella *et al*., 1992; Pijnappels *et al*., 1998; Christensen *et al*., 1999; Hiersemenzel *et al*., 2000; Frigon & Rossignol, 2008; Hurteau & Frigon, 2018; Duysens 2024).

Spinal cord injury (SCI) disrupts ascending and descending pathways that communicate between the brain and spinal networks controlling arm/forelimb and leg/hindlimb movements, including those activated and modulated by cutaneous inputs. The disruption of these pathways and associated sensorimotor deficits in balance and limb coordination vary depending on the severity and level of the lesion (Barbeau *et al*., 2002; Edgerton *et al*., 2004; Frigon & Rossignol, 2006; Rossignol & Frigon, 2011). To investigate how SCI affects cutaneous reflex pathways projecting to the four limbs, we recently used a staggered thoracic lateral hemisections paradigm in cats and evoked cutaneous reflex responses by stimulating the SP nerve (Mari *et al*., 2024). Staggered thoracic hemisections disrupt direct communication between the brain/cervical cord and lumbar locomotor networks, disrupting fore-hind coordination and balance in cats and rats (Jane *et al*., 1964; Kato *et al*., 1984, 1985; Stelzner & Cullen, 1991; Courtine *et al*., 2008; Van Den Brand *et al*., 2012; Cowley *et al*., 2015; Audet *et al*., 2023). We reported a significant reduction in the presence of mid- (19-34 ms) and long-latency (35-60 ms) responses in muscles of all four limbs, especially in the forelimbs (Mari *et al*., 2024). In humans, a few studies have shown changes in interlimb reflexes (i.e. from the arms to the legs or from the legs to the arms) after spinal cord injury, but only at rest (Calancie, 1991; Calancie *et al*., 1996, 2002; Butler *et al*., 2016). Thus, we do not know how reflexes evoked by forelimb afferents are reorganized after SCI during locomotion.

Therefore, the purpose of the present study was to investigate reflex responses evoked by stimulating the SR nerve before and after staggered thoracic hemisections in the same animal during treadmill locomotion, extending our previous findings with responses evoked by hindlimb cutaneous afferents (Mari *et al*., 2024). We first assessed SR-evoked reflex responses in the intact state and then following a first lateral hemisection at mid-thoracic (T5-T6) on the right side. We then assessed how SR-evoked reflex responses changed following a second lateral hemisection at left T10-T11. The main finding was a loss/reduction in homolateral and diagonal responses in hindlimb muscles after staggered thoracic hemisections, which correlated with weakened coordination between the fore- and hindlimbs and impaired balance during quadrupedal locomotion (Audet *et al*., 2023).

## MATERIAL AND METHODS

### Ethical approval

All procedures were approved by the Animal Care Committee of the Université de Sherbrooke (Protocol 442-18) in accordance with policies and directives of the Canadian Council on Animal Care. We obtained the current data set from seven adult purpose-bred cats (> 1 year of age at the time of experimentation), 3 females and 4 males, weighing between 3.4 kg and 6.5 kg, purchased from Marshall BioResources (North Rose, NY, USA). Before and after experiments, cats were housed and fed (weight-dependent metabolic diet and water ad libitum) in a dedicated room within the animal care facility of the Faculty of Medicine and Health Sciences at the Université de Sherbrooke. We followed the ARRIVE guidelines 2.0 for animal studies (Grundy, 2015; Percie Du Sert *et al*., 2020). The investigators understand the ethical principles under which the journal operates and our work complies with this animal ethics checklist. In order to maximize the scientific output of each animal, they were used in other studies to investigate different scientific questions, some of which have been published (Lecomte *et al*., 2022, 2023; Merlet *et al*., 2022; Audet *et al*., 2023; Mari *et al*., 2023, 2024).

### General surgical procedures

All surgeries (implantation and spinal lesions) were performed under aseptic conditions with sterilized equipment in an operating room. Prior to surgery, cats were sedated with an intramuscular (i.m.) injection of butorphanol (0.4 mg/kg), acepromazine (0.1 mg/kg), and glycopyrrolate (0.01 mg/kg). We then injected a mixture (0.05 ml/kg, i.m.) of diazepam (0.25 mg/kg) and ketamine (2.0 mg/kg) in a 1:1 ratio five minutes later for induction. We shaved the animal’s fur (back, stomach, fore- and hindlimbs) and cleaned the skin with chlorhexidine soap. Cats were anesthetized with isoflurane (1.5-3%) and O_2_ delivered with a mask and then with a flexible endotracheal tube. The depth of anesthesia was confirmed by applying pressure to a paw (to detect limb withdrawal) and by assessing the size and reactivity of pupils. Isoflurane concentration was adjusted throughout the surgery by monitoring cardiac and respiratory rates. Body temperature was maintained constant (37 ± 0.5°C) using a water-filled heating pad placed under the animal, an infrared lamp placed ∼50 cm over it and a continuous infusion of lactated Ringers solution (3 ml/kg/h) through a catheter placed in a cephalic vein. At the end of surgery, we injected subcutaneously an antibiotic (cefovecin, 8 mg/kg) and a fast-acting analgesic (buprenorphine, 0.01 mg/kg). We also taped a fentanyl (25 µg/h) patch to the back of the animal 2-3 cm rostral to the base of the tail for prolonged analgesia, which we removed 4-5 days later. After surgery, cats were placed in an incubator and closely monitored until they regained consciousness. We administered another dose of buprenorphine ∼7 hours after surgery. At the end of experiments, cats were anaesthetized with isoflurane (1.5–3.0%) and O2 before receiving a lethal dose of pentobarbital (120 mg/kg) through the left or right cephalic vein. Cardiac arrest was confirmed using a stethoscope to determine the death of the animal. Spinal cords were then harvested for histological analysis (Lecomte *et al*., 2022, 2023; Audet *et al*., 2023; Mari *et al*., 2024).

### Staggered lateral hemisections

After collecting data in the intact state, we performed a lateral hemisection between the 5th and 6th thoracic vertebrae (T5-T6) on the right side of the spinal cord. Before surgery, we sedated the cat with an intramuscular injection of a cocktail containing butorphanol (0.4 mg/kg), acepromazine (0.1 mg/kg) and glycopyrrolate (0.01 mg/kg) and inducted with another intramuscular injection (0.05 ml/kg) of ketamine (2.0 mg/kg) and diazepam (0.25 mg/kg) in a 1:1 ratio. We shaved the fur overlying the back and the skin was cleaned with chlorhexidine soap. The cat was then anesthetized with isoflurane (1.5-3%) and O2 using a mask for a minimum of 5 minutes and then intubated with a flexible endotracheal tube. Isoflurane concentration was confirmed and adjusted throughout the surgery by monitoring cardiac and respiratory rates, by applying pressure to the paw to detect limb withdrawal and by assessing muscle tone. Once the animal was deeply anesthetized, an incision of the skin over and between the 5th and 6^th^ thoracic vertebrae (T5-T6) was made and after carefully setting aside muscle and connective tissue, a small laminectomy of the corresponding dorsal bone was performed. Lidocaine (xylocaine, 2%) was applied topically followed by 2-3 intraspinal injections on the right side of the cord. We then sectioned the spinal cord laterally from the midline to the right using surgical scissors. We placed hemostatic material (Spongostan) within the gap before sewing back muscles and skin in anatomical layers. In the days following hemisection, voluntary bodily functions were carefully monitored. The bladder and large intestine were manually expressed if needed. Once data were collected following the first hemisection (9-13 weeks), we performed a second lateral hemisection between the 10th and 11th thoracic vertebrae (T10-T11) on the left side of the spinal cord using the same surgical procedures and post-operative care described above.

### Electromyography and nerve stimulation

To record the electrical activity of muscles (EMG, electromyography), we directed pairs of Teflon-insulated multistrain fine wires (AS633; Cooner Wire Co., Chatsworth, CA, USA) subcutaneously from two head-mounted 34-pin connectors (Omnetics Connector Corp., Minneapolis, MN, USA). Two wires, stripped of 1–2 mm of insulation, were sewn into the belly of selected forelimb/hindlimb muscles for bipolar recordings. The head-mounted connectors were fixed to the skull using dental acrylic and four to six screws. We verified electrode placement during surgery by stimulating each muscle through the appropriate head connector channel to assess the biomechanically desired muscle contraction. During experiments, EMG signals were pre-amplified (×10, custom-made system), bandpass filtered (30–1,000 Hz) and amplified (100– 5,000×) using a 16-channel amplifier (model 3500; AM Systems, Sequim, WA, USA). EMG data were digitized (5,000 Hz) with a National Instruments (Austin, TX, USA) card (NI 6032E), acquired with custom-made acquisition software and stored on computer. Five forelimb muscles were implanted bilaterally: biceps brachii (BB, elbow and shoulder flexor), extensor carpi ulnaris (ECU, wrist dorsiflexor), flexor carpi ulnaris (FCU, wrist plantarflexor), latissimus dorsi (LD, shoulder retractor), and the long head of the triceps brachii (TRI, elbow and shoulder extensor). Ten hindlimb muscles were implanted bilaterally: biceps femoris anterior (BFA, hip extensor), biceps femoris posterior (BFP, hip extensor and knee flexor), iliopsoas (IP, hip flexor), lateral gastrocnemius (LG, ankle plantarflexor and knee flexor), medial gastrocnemius (MG, ankle plantarflexor and knee flexor), sartorius anterior (SRT, hip flexor and knee extensor), semitendinosus (ST, knee flexor and hip extensor), soleus (SOL, ankle plantarflexor), tibialis anterior (TA, ankle dorsiflexor), and vastus lateralis (VL, knee extensor).

For bipolar nerve stimulation, pairs of Teflon-insulated multistrain fine wires (AS633; Cooner Wire Co., Chatsworth, CA, USA) were passed through a silicon tubing. A horizontal slit was made in the tubing and wires within the tubing were stripped of their insulation. The ends protruding through the cuff were knotted to hold the wires in place and glued. The ends of the wires away from the cuff were inserted into four-pin connectors (Hirose or Samtec) and fixed to the skull using dental acrylic. Cuff electrodes were directed subcutaneously from head-mounted connectors to the left and right SR nerves at the wrist which are purely cutaneous at these levels.

### Experimental design

We collected EMG and kinematic data before and at different time points after staggered hemisections during quadrupedal locomotion at the cat’s preferred treadmill speed (0.3-0.5 m/s). Cats KA and KI stepped at 0.3 and 0.5 m/s, respectively, while the other five cats stepped at 0.4 m/s. The treadmill consisted of two independently controlled belts 130 cm long and 30 cm wide (Bertec) with a Plexiglas separator (130 cm long, 7 cm high, and 1.3 cm wide) placed between the two belts to prevent limbs impeding each other. In the intact, preoperative state, cats were trained for 2-3 weeks in a progressive manner, first for a few steps and then for several consecutive minutes, using food and affection as rewards. Once cats could perform 3-4 consecutive minutes, we started the experiments. During experiments, we delivered trains of electrical stimuli consisting of three 0.2 ms pulses at 300 Hz using a Grass (West Warwick, RI, USA) S88 stimulator. At the start of the experiment, we determined the motor threshold, defined as the minimal intensity that elicited a small motor response in an ipsilateral flexor muscle (e.g., ST or TA) during the swing phase. We then set stimulation intensity at 1.2 times the motor threshold to mainly activate large diameter Aβ cutaneous afferents. A locomotor trial lasted 4-5 min and consisted of ∼60 stimuli delivered pseudo-randomly every 2–4 cycles. Stimuli were delivered at specific points of the stimulated forelimb (left or right) movement: mid-stance, the transition from stance-to-swing, mid-swing and the transition from swing-to-stance. At the start of the experiment, stimulation delays were determined using real-time EMG to detect the onset of an extensor burst in relation to stance and swing phases. We then set delays in relation to this extensor EMG so that stimuli were delivered at the four desired time points. The timing of the stimuli was assessed during off-line analysis and stimuli that did not fall in the desired phases were excluded. We characterized responses in muscles of the stimulated forelimb (homonymous), the opposite forelimb (crossed), the hindlimb on the same side (homolateral) and the diagonal hindlimb (diagonal). Figure 1A describes the timeline of data collection in all seven cats at two time points after the first hemisection (**H1T1** and **H1T2** in 7 cats) and/or at 1-2 time points after the second hemisection (**H2T1** in 2 cats and **H2T2** in 6 cats). Some cats only have one time point after the second hemisection because they took longer to recover quadrupedal locomotion. No data were collected for cat KI after the second hemisection due to technical issues with the implants.

**Figure 1.**
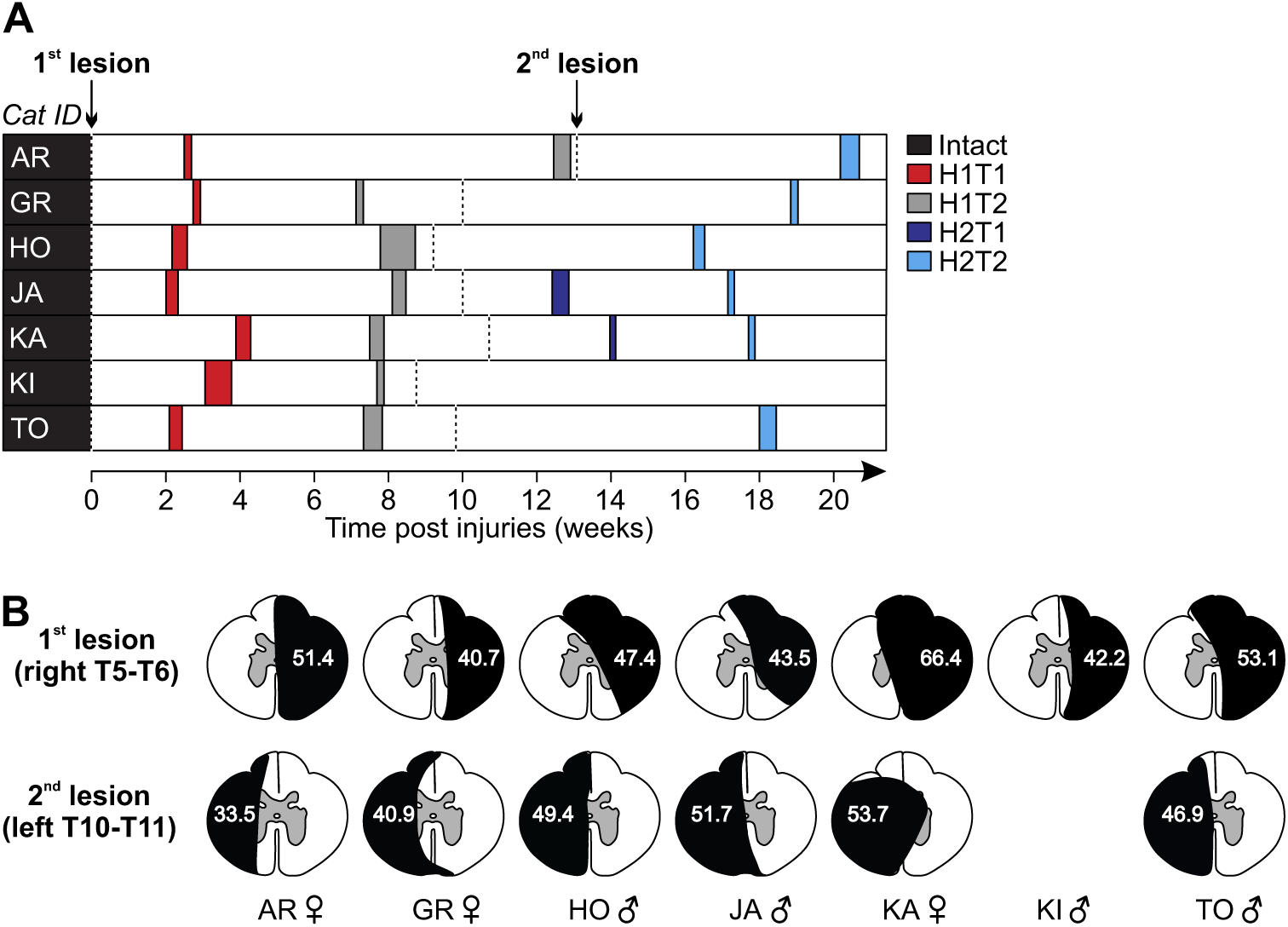
Experimental chronology and estimation of lesions extent. **(A)** Chronology showing the first (T1) and second (T2) experimental time points after the first (H1) and second (H2) hemisections in all cats. **(B)** For the extent of the lesions, the black area represents the estimation as a percentage of total cross-sectional area (reproduced with permission of (Mari *et al*., 2024)). Note that we only performed one lesion in cat KI.

### Histology

After confirming euthanasia (i.e., no cardiac and respiratory functions), we harvested an approximately 2 cm long section of the spinal cord centered on the lesions. Segments of the dissected spinal cord were then placed in a 25 ml 4% paraformaldehyde (PFA) solution (in 0.1 M phosphate-buffered saline (PBS), 4°C). After 5 days, we placed the spinal cord in a new PBS solution containing 30% sucrose for 72 h at 4°C, then froze it in isopentane at −50°C for cryoprotection. The spinal cord was then sliced in 50 µm coronal sections using a cryostat (Leica CM1860, Leica BioSystems Inc., Concord, ON, Canada) and mounted on gelatinized-coated slides. The slides were dried overnight and then stained with a 1% cresyl violet acetate solution for 12 min. We washed the slides for 3 min in distilled water before being dehydrated in successive baths of ethanol (50%, 70% and 100%, 5 min each) and transferring them in xylene for 5 min. Dibutylphthalate polystyrene xylene was next used to mount and dry the spinal cord slides before being scanned by a Nanozoomer 2.0-RS (Hamamatsu Corp., Bridgewater, NJ, USA). We then performed qualitative and quantitative analyses to estimate lesion extent using ImageJ by selecting the slide with the greatest identifiable damaged area. Using the scarring tissue stained with cresyl violet acetate, we estimated lesion extent by dividing the lesion area by the total area of the selected slice and expressed it as percentage. Lesion extent estimations for individual cats after the first and second spinal lesions are shown in Figure 1B. They ranged from 40.7% to 66.4% (49.2 ± 8.9%) and 33.5% to 53.7% (46.0 ± 7.6%) for the first and second hemisections, respectively.

### Reflex analysis

We described the reflex analysis in several of our publications (Hurteau *et al*., 2017, 2018; Hurteau & Frigon, 2018; Merlet *et al*., 2020, 2021; Mari *et al*., 2023), and recently illustrated it in detail in (Mari *et al*., 2024). Briefly, for all locomotor sessions, EMG signals were low-pass filtered (250 Hz) to facilitate the visualization of the EMG activity envelope. We first defined locomotor cycles from successive burst onsets of an extensor from the stimulated forelimb, then separated them as stimulated (i.e., cycles with stimulation) or control (i.e., cycles without stimulation) cycles. Sections where the cat stepped irregularly were removed from analysis based on EMG and video data. Stimulated cycles were then sorted and divided into 4 subphases based on stance onset of the stimulated forelimb: swing-to-stance, mid-stance, stance-to-swing and mid-swing. Control (^C̅^) cycles were averaged and rectified to provide a baseline locomotor EMG, an indication of the excitability level of the motor pool at stimulation. We averaged the stimulated (^S̅^) cycles and time normalized ^C̅^ to ^S̅^ cycle durations and superimposed them. To determine response onsets and offsets, defined as prominent positive or negative deflections away from ^C̅^, we set windows using previous studies as guidelines (Duysens & Stein, 1978; Duysens & Loeb, 1980; Pratt *et al*., 1991; Loeb, 1993; Hurteau *et al*., 2017, 2018; Hurteau & Frigon, 2018; Mari *et al*., 2023) with 97.5% confidence intervals. We termed short-latency (7–18 ms; **SLR**) excitatory and inhibitory responses as P1 and N1 responses, respectively, based on the terminology introduced by (Duysens & Loeb, 1980). Responses in the crossed, homolateral, and diagonal limbs that had an onset ≤18 ms were classified as P1 or N1, as the minimal latency for spino-bulbo-spinal reflexes in the cat is 16-18 ms (Shimamura & Livingston, 1962; Shimamura *et al*., 1990). Mid-latency (19–34 ms; **MLR**) excitatory and inhibitory responses were termed P2 and N2, respectively. Long-latency (35–60 ms; **LLR**) excitatory and inhibitory responses were termed P3 and N3, respectively. The EMG of reflex responses ^S̅^ was then integrated and subtracted from the integrated ^C̅^ in the same time window to provide a net reflex value. This net reflex value was then divided by the integrated ^C̅^ value to evaluate reflex responses. This division helps identify if changes in reflex responses across the cycle are independent of changes in ^C̅^ activity (Matthews, 1986; Frigon & Rossignol, 2007, 2008, 2009; Hurteau *et al*., 2017, 2018; Hurteau & Frigon, 2018; Mari *et al*., 2023, 2024).

### Statistical analysis

We performed statistical tests with IBM SPSS Statistics V26 (IBM Corp., Armonk, NY, USA). We quantified reflex responses in five forelimb muscles (BB, ECU, FCU, LD, and TRI) and in ten hindlimb muscles (BFA, BFP, IP, LG, MG, SRT, SOL, ST, TA, and VL) when stimulation was delivered to the left or right SR. To evaluate whether homonymous, crossed, homolateral and diagonal responses were modulated by phase, we performed a one factor (phase) ANOVA on all responses (P1, P2, P3, N1, N2 and N3) in each cat and state/time point. Because we have several responses within a given phase, we considered all responses during a locomotor session as a population. In our statistical analysis, we used mixed models to deal with incomplete data sets. For instance, reflex responses are sometimes absent after spinal lesions. Response occurrence probabilities, defined as the fraction of evoked responses obtained out of all cats for pooled SLR, MLR and LLR from the different states/time points were compared using a generalized linear mixed model (GLMM) with a binomial distribution and a logit link (mixed logistic regression) in all four limbs. The GLMM analysis was performed using state/time point as a fixed factor. We incorporated random intercepts at two distinct levels to consider the hierarchical relationships present in our dataset. A random intercept on individual cats at the upper level captured variability across cats. A random intercept on muscle nested within cat at a lower level, acknowledging that the same muscle response data were repeatedly measured within each cat to help us account for any correlation or non-independence of observations within the same cat-muscle pair. Statistical significance for all tests was set at p < 0.05.

## RESULTS

### Recovery of quadrupedal locomotion and changes in fore-hind coordination after staggered hemisections

We recently described changes in the quadrupedal locomotor pattern after staggered thoracic lateral hemisections (right T5-T6 followed by left T10-T11), including six cats of the present study (Audet *et al*., 2023). Briefly, we showed that cats spontaneously recovered quadrupedal locomotion following both hemisections but required balance assistance after the second one. After the first and second hemisections, the coordination between the forelimbs and hindlimbs became weaker and displayed 2:1 patterns, where the forelimbs performed two cycles within one hindlimb cycle.

After the first hemisection (H1), all seven cats regained quadrupedal locomotion on the treadmill within one to two weeks. We were able to conduct reflex sessions for several consecutive minutes at the first (H1T1) and second (H1T2) time points. After the second hemisection (H2), the six cats tested recovered quadrupedal locomotion within two to five weeks. However, they required mediolateral balance assistance during reflex sessions, which was provided by an experimenter holding the tail of the animal but without providing weight support. As stated in the Methods, some cats only have one time point after the second hemisection due to the longer recovery of quadrupedal locomotion and their ability to maintain it for reflex testing. Only two cats, KA and JA, participated in reflex sessions at the first time point (H2T1), i.e., approximately two weeks after the second hemisection. All six cats performed reflex sessions eight weeks later at the second time point (H2T2).

### Cutaneous reflexes evoked by stimulating the superficial radial nerve before and after staggered hemisections

To assess the reorganization of cutaneous reflex pathways from forelimb afferents after SCI, we stimulated the left and right SR nerves before and after the first and second hemisections and recorded reflex responses during quadrupedal treadmill locomotion in muscles of the four limbs. Figures 2-5 illustrate examples from representative cats for homonymous, crossed, homolateral and diagonal responses in selected muscles at four phases of the cycle (swing-to-stance transition, mid-stance, stance-to-swing transition and mid-swing). We observed several changes in reflex responses after the first and second hemisections, but for the responses shown in Figures 2-5, we only highlight the most noticeable observations. **Tables 1-3** provide the full details of reflex responses before and after staggered hemisections for individual cats. For each state/time point, filled areas represent evoked responses and are optimized for display according to the strongest response obtained in one of the four phases. If an area is not filled, it means that the stimulated EMG did not deviate sufficiently from the baseline EMG to be defined as a response. The scale is optimized per state/time point and differs across state/time points. This is to show the pattern of evoked responses and its phase-dependent modulation at a given state/time point, as in our recent study (Mari *et al*., 2024)

**Figure 2.**
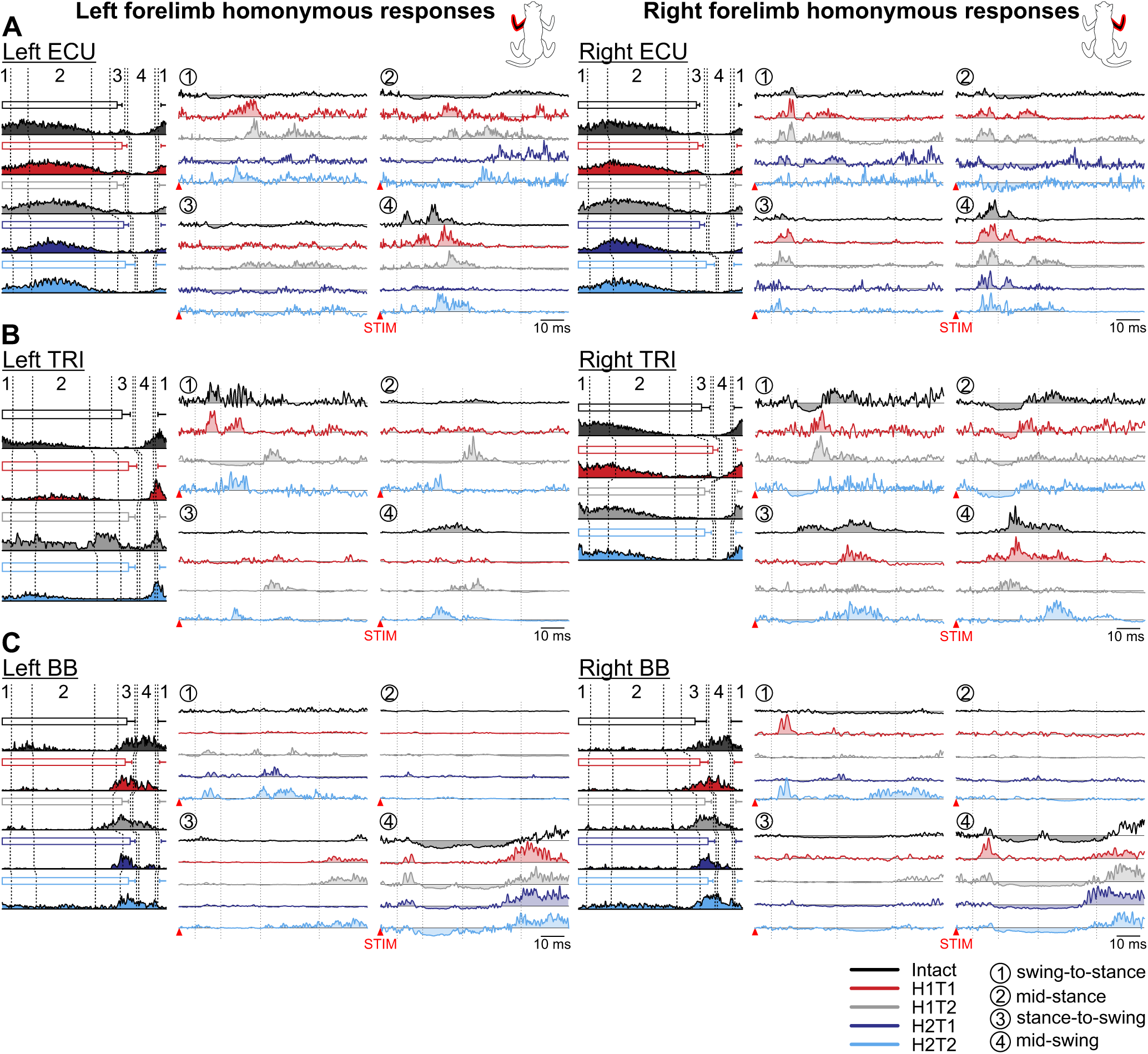
Phase-dependent modulation of cutaneous reflexes evoked in homonymous forelimb muscles during locomotion before and following staggered hemisections. Each panel shows, from left to right, stance phases of the stimulated forelimb (empty horizontal bars) with its averaged rectified muscle activity normalized to cycle duration in the different states/time points, and homonymous reflex responses in representative cats for the left and right **(A)** extensor carpi ulnaris (ECU, cat KA), **(B)** triceps brachii (TRI, cat TO), and **(C)** biceps brachii (BB, cat JA). Reflex responses are shown with a post-stimulation window of 80 ms in four phases in the intact state, and after the first (H1) and second (H2) hemisections at time points 1 (T1) and/or 2 (T2). At each state/time point, evoked responses are scaled according to the largest response obtained in one of the four phases. The scale, however, differs between states/time points. The dotted vertical lines in the reflex responses indicate the N1/P1, N2/P2 and N3/P3 time windows.

**Table 1.**
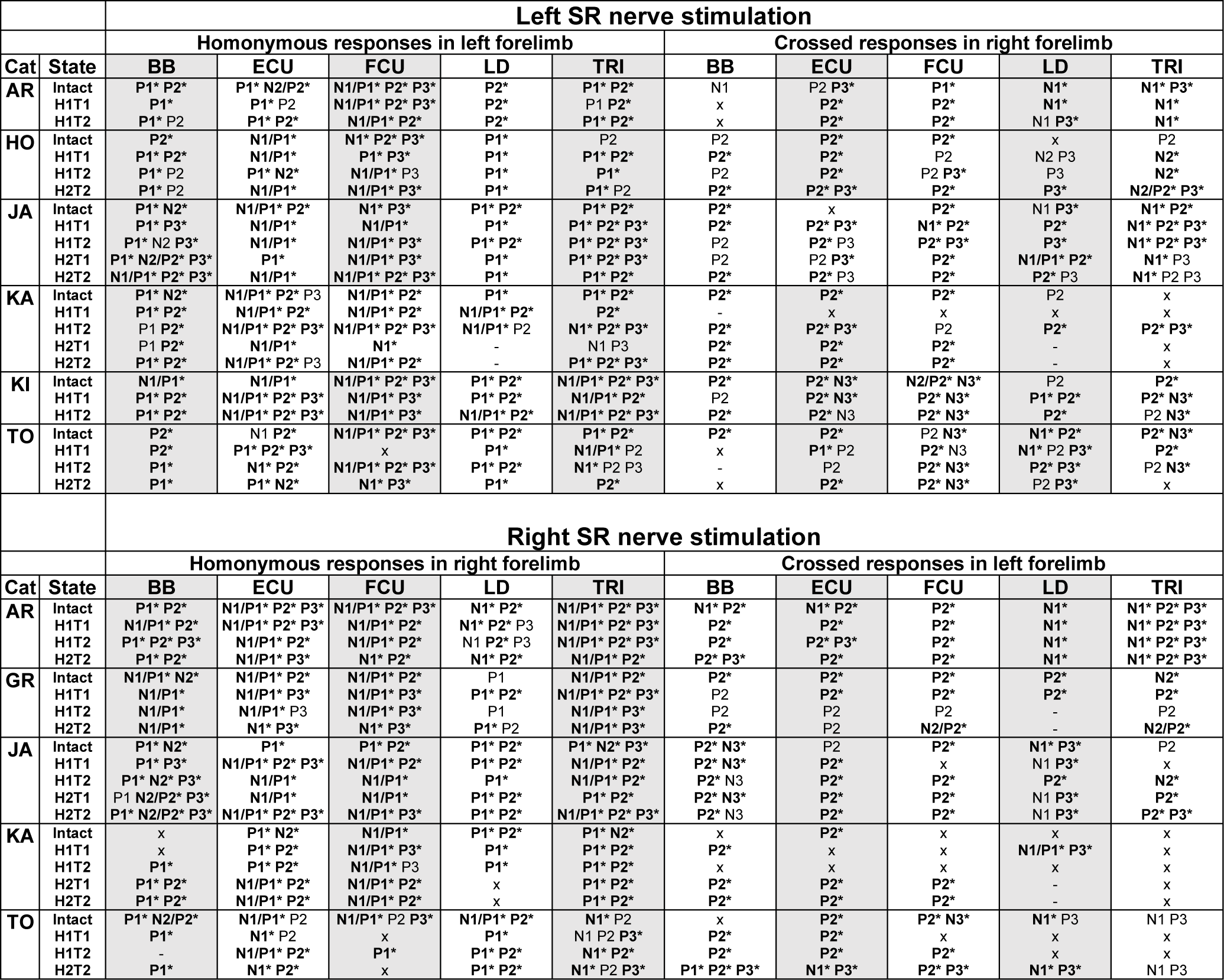
Homonymous and crossed reflex response before and after staggered hemisections. The table shows homonymous and crossed responses (P1, P2, P3, N1, N2 and N3) evoked in individual cats in left and right forelimb muscles in the intact state, and after the first (H1) and second (H2) hemisections at 1-2 time points (T1 or T2). Bold responses with an asterisk indicate a significant phase modulation (one factor ANOVA, p < 0.05). x, No response. -, Non-implanted or non-analyzable (lost or excessive noise) muscle. BB, biceps brachii; ECU, extensor carpi ulnaris; FCU, flexor carpi ulnaris; LD, latissimus dorsi; TRI, triceps brachii.

#### Homonymous responses in forelimb muscles

We stimulated the left and right SR nerves and recorded homonymous responses in muscles of the left and right forelimbs, respectively, before and after staggered hemisections (Fig. 2 and **Table 1**). We illustrate homonymous reflex responses in three muscles (ECU, TRI and BB) bilaterally in representative cats. The ECU and TRI muscles are mostly active during stance while BB is active during swing and/or at the stance-to-swing transition. We observed that the burst profiles of the three selected forelimb muscles remained similar across states (before and after hemisections) and time points during the locomotor cycle.

In the left ECU (Fig. 2A, left panel), in the intact state, we observed homonymous P1/P2 responses at mid-swing, and weak N1 responses in the other three phases followed by small P3 responses at mid-stance and stance-to-swing. After the first hemisection, at H1T1 and H1T2, P1/P2 responses remained at mid-swing, but relatively strong P2 and/or P3 responses appeared at swing-to-stance and mid-stance. After the second hemisection, at H2T1, P2/P3 responses were reduced at mid-swing and absent in the other phases. At H2T2, P1/P2 responses recovered at mid-swing and returned at swing-to-stance and mid-stance. In the right ECU (Fig. 2A, right panel), in the intact state, we observed P1 responses in all phases followed by N2 at swing-to-stance and mid-stance and P2 at mid-swing. At H1T1 and H1T2, P1 responses were prominent in all phases followed by prominent P2 and/or P3 responses, except for stance-to-swing with only P1 at H1T2. At H2T1, P1/P2 responses remained at swing-to-stance and mid-swing, with N1 at mid-stance and no responses at stance-to-swing. At H2T2, we observed P1 responses at mid-swing and stance-to-swing, and N1 responses at mid-stance, with no P2/P3 responses in all phases.

In the left TRI (Fig. 2B, left panel), we observed homonymous P1 and P2 responses at swing-to-stance and P2 responses at mid-swing and mid-stance. At H1T1, P1/P2 and P2 responses remained at swing-to-stance and mid-stance, respectively, but P2 responses were lost at mid-swing. At H1T2, we N1 responses followed by P3 responses appeared at swing-to-stance and mid-stance. At H2T2, we only observed P2 responses in all phases. In the right TRI (Fig. 2B, right panel), we observed P2/P3 responses in all phases that followed N1 responses at swing-to-stance and mid-stance. At H1T1, N1 responses were lost or weakened at swing-to-stance and mid-stance, respectively, but P2 responses remained. N2 responses appeared at stance-to-swing and P1 responses at mid-swing, while P2/P3 remained in both phases. At H2T1, a weak N1/P2 responses returned or remained stance-to-swing and mid-stance but excitatory responses were lost or reduced at stance-to-swing and mid-swing. At H2T2, N1 responses were prominent at swing-to-stance and mid-stance, but P2 responses were lost or reduced. At stance-to-swing and mid-swing prominent P2/P3 returned.

In the left BB (Fig. 2C, left panel), the most noticeable response was a prolonged N1 response at mid-swing in the intact state. At H1T1, this N1 response was lost and a prominent P3 response appeared. At H1T2, the N1 response returned followed by a P3 response. These N1/P3 responses remained at H2T1 and H2T2. The other noticeable change was at swing-to-stance, with the appearance of P1 responses at H1T2, which remained at H2T1 and H2T2. At H2T1 and H2T2, P2/P3 responses also appeared. In the right BB (Fig. 2C, right panel), in the intact state, the most noticeable response at mid-swing was an N2 response that followed a brief P1 response. At H1T1, this N2 response was lost while the P1 response became prominent. The N2 response returned at H1T2 following a brief P1. At H2T1 and H2T2, the N2 response was prominent and was followed by prominent P3 responses. In the other phases, the only noticeable changes were the appearance of a prominent P1 responses at H1T1 at swing-to-stance, which disappeared at H1T2 before returning at H2T2.

To summarize, even though thoracic hemisections did not directly affect the pathways transmitting cutaneous afferent inputs from the SR nerve to ipsilateral (homonymous) motor circuits in the cervical cord, we observed several small reflex changes in forelimb muscles.

#### Crossed responses in forelimb muscles

We stimulated the left and right SR nerves and recorded crossed responses in muscles of the right and left forelimbs, respectively, before and after staggered hemisections in the four phases (Fig. 3 and **Table 2**). We defined the four phases according to the stimulated limb, but it is important to consider the phase of the contralateral limb where the responses were recorded.

**Figure 3.**
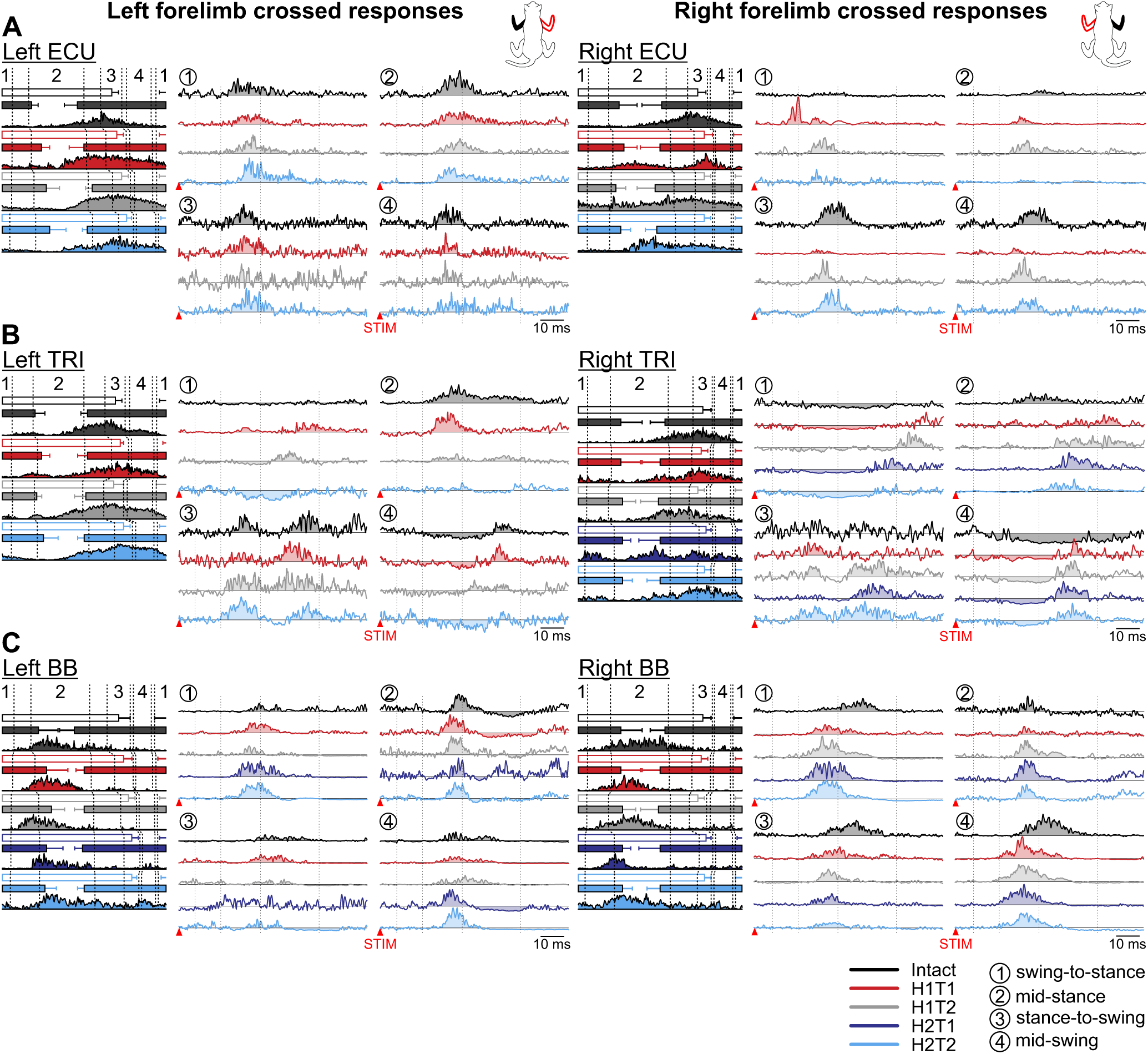
Phase-dependent modulation of cutaneous reflexes evoked in crossed forelimb muscles during locomotion before and following staggered hemisections. Each panel shows, from left to right, stance phases of the stimulated forelimb (empty horizontal bars) and crossed forelimb (filled horizontal bars) with its averaged rectified muscle activity normalized to cycle duration in the different states/time points, and crossed reflex responses in representative cats for the left and right **(A)** extensor carpi ulnaris (LECU, cat GR; RECU, cat TO), **(B)** triceps brachii (LTRI, cat AR; RTRI, cat JA), and **(C)** biceps brachii (BB, cat JA). Reflex responses are shown with a post-stimulation window of 80 ms in four phases in the intact state, and after the first (H1) and second (H2) hemisections at time points 1 (T1) and/or 2 (T2). At each state/time point, evoked responses are scaled according to the largest response obtained in one of the four phases. The scale, however, differs between states/time points. The dotted vertical lines in the reflex responses indicate the N1/P1, N2/P2 and N3/P3 time windows.

**Table 2.**
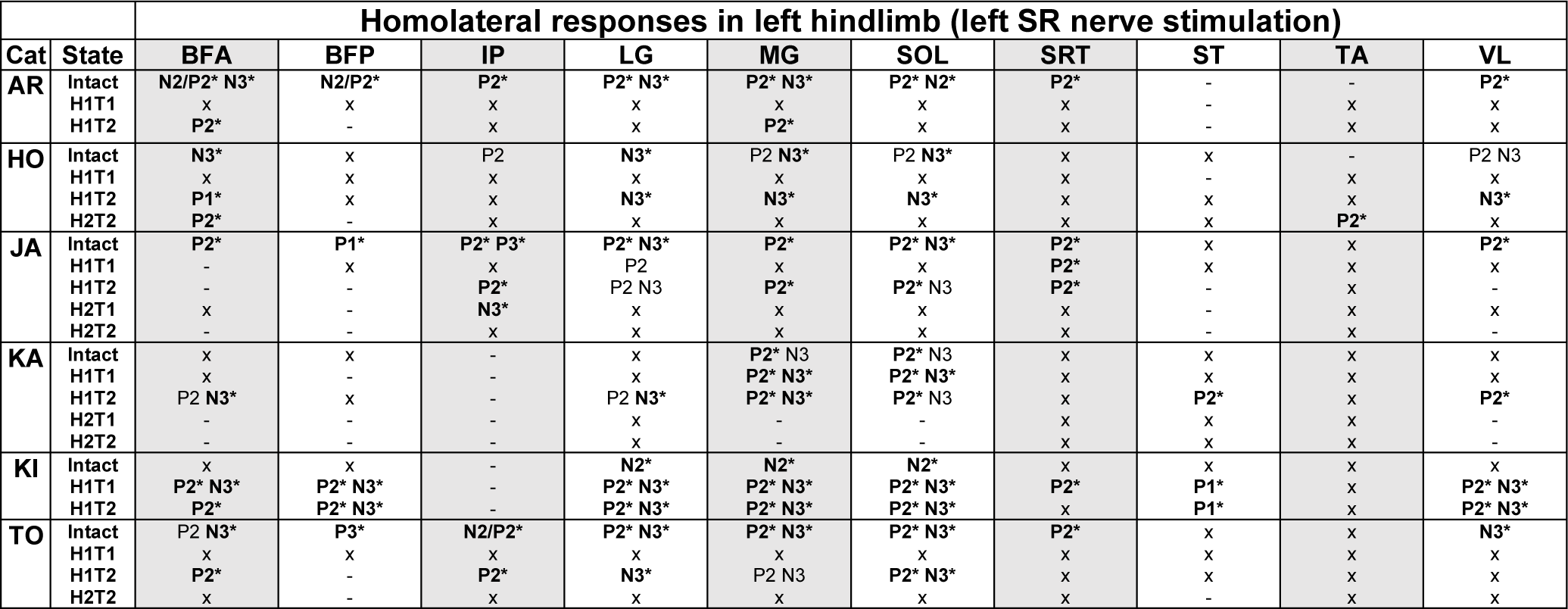
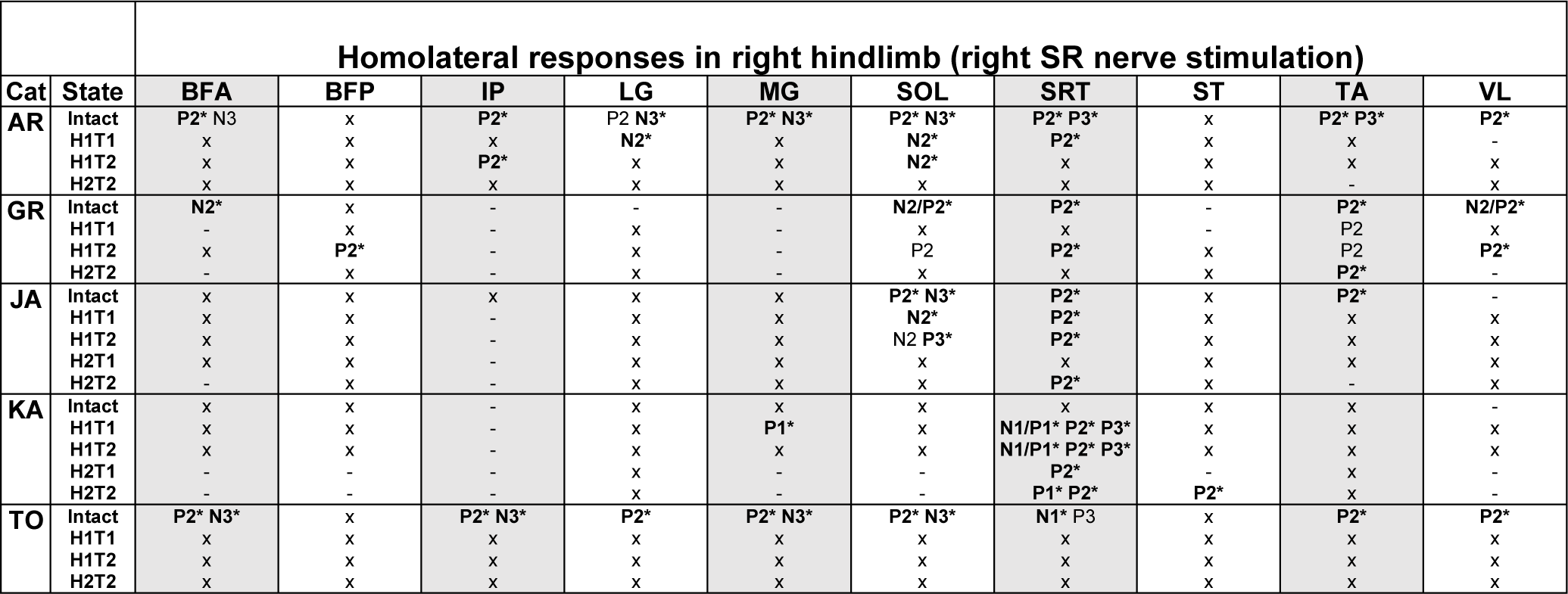
Homolateral reflex responses before and after staggered hemisections. The table shows homolateral responses (P1, P2, P3, N1, N2 and N3) evoked in individual cats in left and right hindlimb muscles in the intact state, and after the first (H1) and second (H2) hemisections at 1-2 time points (T1 or T2). Bold responses with an asterisk indicate a significant phase modulation (one factor ANOVA, p < 0.05). x, No response. -, Non-implanted or non-analyzable (lost or excessive noise) muscle. BFA, biceps femoris anterior; BFP, biceps femoris posterior; IP, iliopsoas; LG, lateral gastrocnemius; MG, medial gastrocnemius; SOL, soleus; SRT, anterior sartorius; ST, semitendinosus; TA, tibialis anterior; VL, vastus lateralis.

**Table 3.**
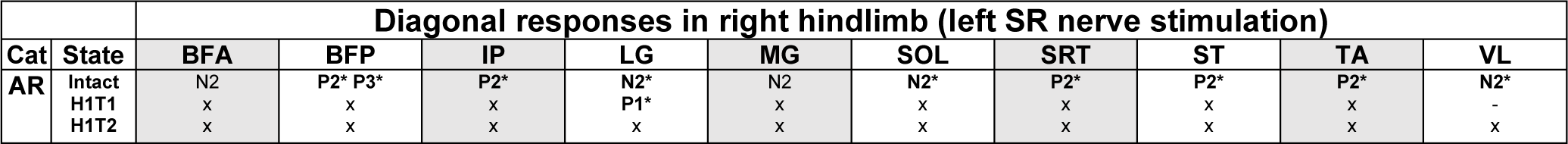
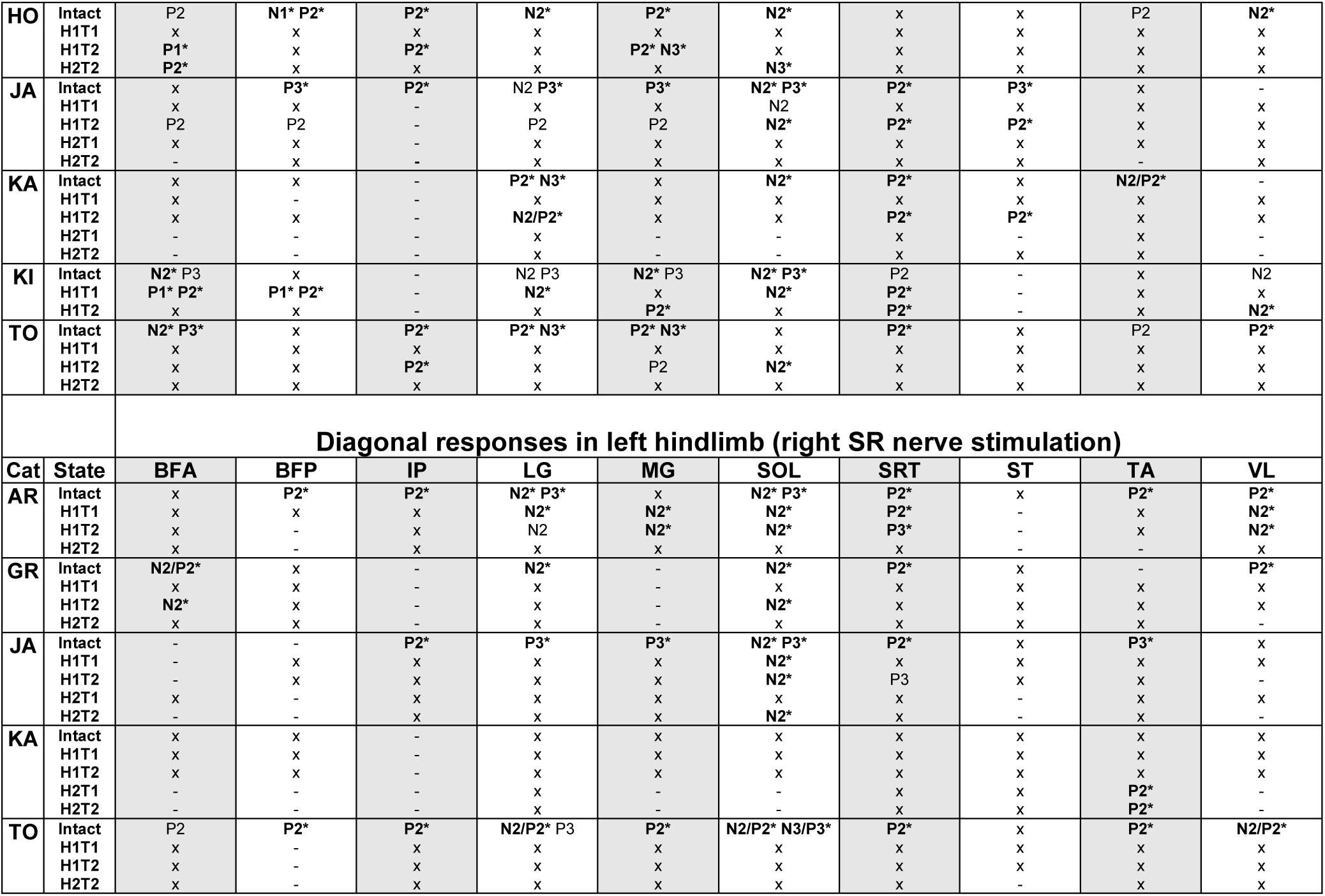
Diagonal reflex responses before and after staggered hemisections. The table shows diagonal responses (P1, P2, P3, N1, N2 and N3) evoked in individual cats in left and right hindlimb muscles in the intact state, and after the first (H1) and second (H2) hemisections at 1-2 time points (T1 or T2). Bold responses with an asterisk indicate a significant phase modulation (one factor ANOVA, p < 0.05). x, No response. -, Non-implanted or non-analyzable (lost or excessive noise) muscle. BFA, biceps femoris anterior; BFP, biceps femoris posterior; IP, iliopsoas; LG, lateral gastrocnemius; MG, medial gastrocnemius; SOL, soleus; SRT, anterior sartorius; ST, semitendinosus; TA, tibialis anterior; VL, vastus lateralis.

In the left ECU (Fig. 3A, left panel), we observed crossed P2 responses in all four phases in the intact state, with the largest at mid-stance of the stimulated right forelimb. After the first and second hemisections, P2 responses remained in all phases. In the right ECU (Fig. 3A, right panel), in the intact state, we observed crossed P2 responses in all phases, with the largest at stance-to-swing and mid-swing of the stimulated left forelimb. After the first hemisection, P2 responses were reduced at stance-to-swing and mid-swing and P1 responses appeared at swing-to-stance. P2 responses recovered at H1T2 and were observed in all phases. P2 responses persisted after the second hemisection at H2T2, except at mid-stance where they were absent.

In the left TRI (Fig. 3B, left panel), in the intact state, we observed crossed N1 responses during mid-swing followed by P3 responses, whereas at mid-stance and stance-to-swing, we observed P2/P3 responses. After the first hemisection, at H1T1 and H1T2, the response pattern was largely maintained, but P2/P3 responses appeared at swing-to-stance at H1T1 and N2/P3 responses at H1T2. After the second hemisection, at H2T2, we observed N2 responses at swing-to-stance and mid-swing, maintained P2/P3 responses at stance-to-swing and a weak P2/P3 response at mid-stance. In the right TRI (Fig. 3B, right panel), in the intact state, we observed prolonged crossed N1 responses at mid-swing and swing-to-stance and a P2 response at mid-stance. After the first and second hemisections, we observed prolonged N1 responses at swing-to-stance followed by P3 responses, but only at H1T2 and H2T1. At mid-stance, we observed P2/P3 responses at H1T1 and then only P3 responses at H1T2, H2T1 and H2T2. At stance-to-swing, we observed P2 responses at H1T1, P2/P3 at H1T2, P3 at H2T1 and P2/P3 at H2T2. At mid-swing, N1 or N2 responses were followed by P3 responses at all time points.

In the left BB (Fig. 3C, left panel), in the intact state, we observed crossed P2/P3 responses in all four phases that peaked at mid-stance of the stimulated right forelimb, which was followed by N3 responses at mid-stance only. After the first and second hemisection, P2/P3 responses were maintained in all phases except at stance-to-swing where they were present at H1T1 and H1T2 but disappeared after the second hemisection. In the right BB (Fig. 3C, right panel), in the intact state, we observed crossed P2/P3 responses in all phases that peaked at mid-swing of the stimulated left forelimb. After the first and second hemisection, P2/P3 responses were maintained in all phases with the strongest responses observed at mid-swing and swing-to-stance.

To summarize, although we noted some changes in crossed reflex responses in forelimb muscles after staggered hemisections, they were mostly similar to those observed in the intact state.

**Table 1** summarizes homonymous and crossed reflex response patterns observed in all 5 forelimb muscles bilaterally in 6 and 5 cats for the left and right SR nerve stimulations, respectively, before and after staggered hemisections. Overall, the reflex response patterns that we observed in forelimb muscles remained generally similar after hemisections, as did their phase-dependent modulation.

#### Homolateral responses in hindlimb muscles

Out of 10 hindlimb muscles, we show examples from three selected muscles, SOL, VL and SRT, bilaterally in representative cats. The SOL and VL muscles are mostly active during stance while SRT is active during swing and/or at the stance-to-swing transition. We define the phases relative to the stimulated forelimb. However, after the first and/or second hemisections, cats frequently performed two forelimb cycles within one hindlimb cycle (i.e. 2:1 fore-hind patterns). This means, for example, that stimulation at mid-stance of one forelimb can elicit reflex response at different phases for the hindlimb (see Discussion).

In the left SOL (Fig. 4A, left panel), in the intact state, we observed homolateral P2 responses at swing-to-stance and mid-swing of the stimulated forelimb that were followed by N3 responses at swing-to-stance and mid-stance. After the first hemisection, we observed no responses at H1T1, but some recovery at H1T2, with P2/N3 responses at swing-to-stance and mid-stance and P2 responses at mid-swing. Homolateral responses were lost after the second hemisection. In the right SOL (Fig. 4A, right panel), homolateral response patterns were similar in the intact state, with P2/N3 at swing-to-stance and mid-swing and N3 at mid-stance. After the first and second hemisections, we observed no homolateral responses.

**Figure 4.**
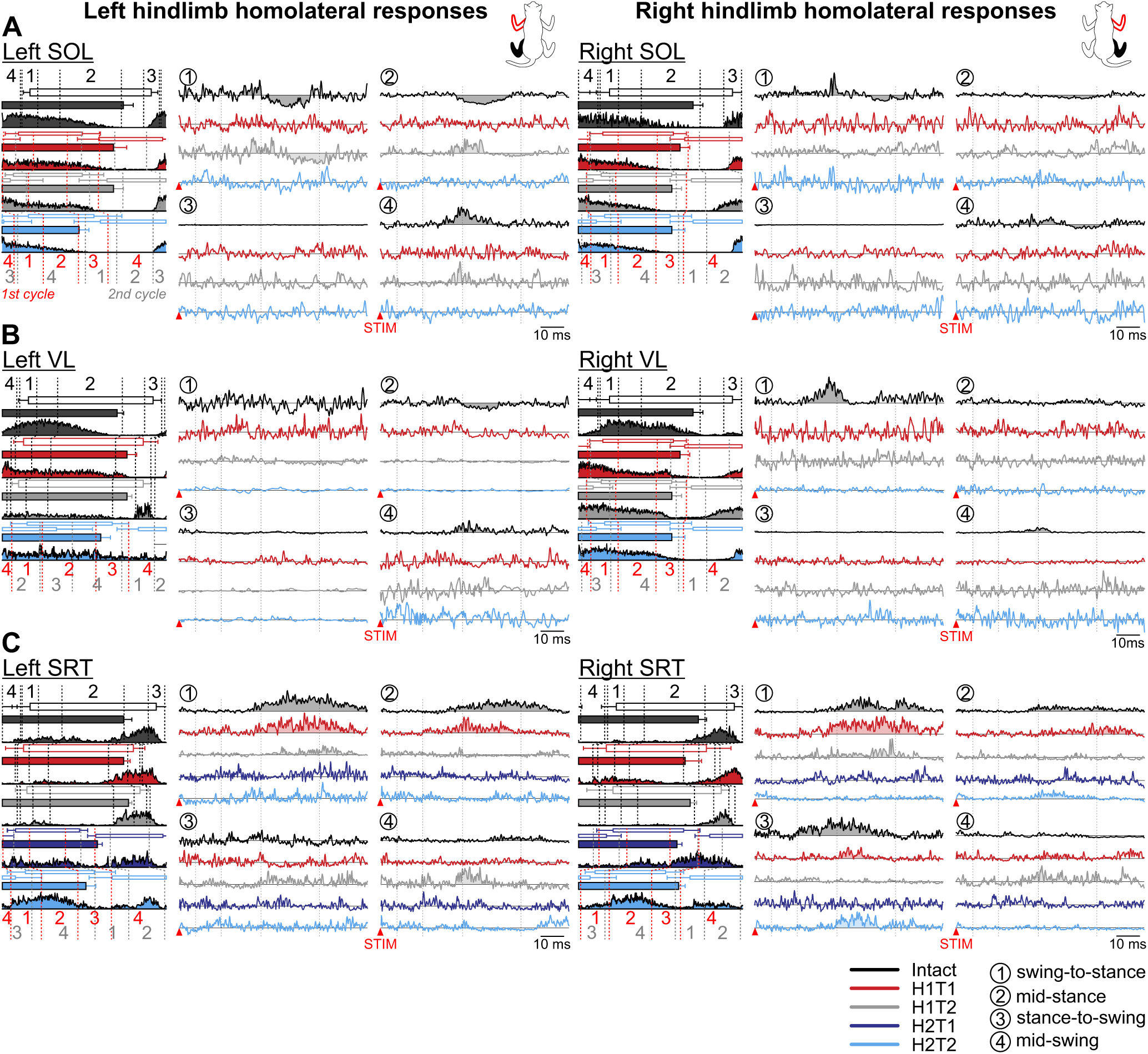
Phase-dependent modulation of cutaneous reflexes evoked in homolateral hindlimb muscles during locomotion before and following staggered hemisections. Each panel shows, from left to right, stance phases of the stimulated forelimb (empty horizontal bars) and homolateral hindlimb (filled horizontal bars) with its averaged rectified muscle activity normalized to cycle duration in the different states/time points, and homolateral reflex responses in representative cats for the left and right **(A)** soleus (SOL, cat TO), **(B)** vastus lateralis (LVL, cat HO; RVL, cat TO), and **(C)** anterior sartorius (SRT, cat JA). Reflex responses are shown with a post-stimulation window of 80 ms in four phases in the intact state, and after the first (H1) and second (H2) hemisections at time points 1 (T1) and/or 2 (T2). At each state/time point, evoked responses are scaled according to the largest response obtained in one of the four phases. The scale, however, differs between states/time points. The dotted vertical lines in the reflex responses indicate the N1/P1, N2/P2 and N3/P3 time windows.

In the left VL (Fig. 4B, left panel), in the intact state, we observed homolateral P2 responses at mid-swing of the stimulated forelimb and N3 responses at mid-stance. After the first and second hemisections, these responses disappeared, with the exception of a weak N3 response at H1T2 swing-to-stance. In the right VL (Fig. 4B, right panel), we observed homolateral P2 responses at swing-to-stance and mid-swing of the stimulated forelimb. After the first and second hemisections, we observed no responses.

In the left SRT (Fig. 4C, left panel), in the intact state, we observed homolateral P2/P3 responses at swing-to-stance, mid-stance and mid-swing of the stimulated left forelimb. After the first hemisection, P2/P3 responses remained at swing-to-stance and mid-stance at H1T1 but were visibly reduced at H1T2. After the second hemisection, we observed no responses. In the right SRT (Fig. 4C, right panel), in the intact state, we observed homolateral P2/P3 responses at swing-to-stance, mid-stance and stance-to-swing of the stimulated right forelimb. After the first hemisection, P2/P3 responses remained in these phases at H1T1 but were visibly reduced or disappeared at H1T2. After the second hemisection, we observed no responses at H2T1 but P2/P3 responses returned at mid-stance and stance-to-swing at H2T2.

**Table 2** summarizes homolateral reflex response patterns in all 10 hindlimb muscles bilaterally in 6 and 5 cats for the left and right SR nerve stimulations, respectively, before and after staggered hemisections. Response patterns mostly consisted of P2 responses followed by N3 responses in extensors (BFA, LG, MG and SOL), and P2/P3 responses in flexors (IP, SRT, ST and TA). Although the phase-dependent modulation of responses generally remained after staggered hemisections, when responses were present, we observed a loss in response occurrence in most muscles after the first and/or second hemisections. However, some homolateral responses returned at H1T2 on the left side.

#### Diagonal responses in hindlimb muscles

In the left SOL (Fig. 5A, left panel), in the intact state, we observed diagonal N2 responses followed by P3 responses at stance-to-swing and mid-swing of the stimulated right forelimb, with P2/N3 responses at mid-stance. After the first and second hemisections, we observed no diagonal responses. In the right SOL (Fig. 5A, right panel), in the intact state, we observed diagonal N2 responses at mid-stance, stance-to-swing and mid-swing of the stimulated left forelimb followed by P3 responses at mid-stance and stance-to-swing. After the first hemisection, at H1T1 and H1T2, N2 responses remained at in these three phases but P3 responses disappeared. After the second hemisection, we observed no diagonal responses.

In the left VL (Fig. 5B, left panel), in the intact state, we observed diagonal P2 responses at swing-to-stance and mid-stance of the stimulated right forelimb, with N2 responses at mid-swing. After the first and second hemisections, we observed no diagonal responses. In the right VL (Fig. 5B, right panel), in the intact state, we observed diagonal P2 responses only at mid-stance of the stimulated left forelimb. After the first and second hemisections, we observed no diagonal responses.

In the left SRT (Fig. 5C, left panel), in the intact state, we observed diagonal P2 responses in all phases except at stance-to-swing of the stimulated right forelimb. After the first and second hemisections, we observed no diagonal responses. In the right SRT (Fig. 5C, right panel), in the intact state, we observed diagonal P2 responses in all phases except at swing-to-stance of the stimulated left forelimb. After the first hemisection, we observed no responses at H1T1 but P2/P3 reappeared at H1T2 at swing-to-stance and mid-stance. After the second hemisection, we observed no diagonal responses.

**Table 3** summarizes diagonal reflex response patterns in all 10 hindlimb muscles bilaterally in 6 and 5 cats for the left and right SR nerve stimulations, respectively, before and after staggered hemisections.

Response patterns consisted mostly of N2 followed by P3 responses in extensors (LG and SOL). In flexors (BFP, IP, SRT, ST and TA), we observed P2/P3 responses. Similar to homolateral responses, we observed a loss in response occurrence in most muscles after the first and/or second hemisections, although the phase-dependent modulation remained if responses were present. We observed some return of diagonal responses at the second time point after the first hemisection (H1T2) in right hindlimb muscles.

#### Staggered hemisections reduce the occurrence of mid- and long-latency responses in hindlimb muscles

After complete or incomplete spinal lesions in cats, mid- and long-latency responses in hindlimb muscles are generally reduced or abolished (Fuwa *et al*., 1991; LaBella *et al*., 1992; Frigon & Rossignol, 2008; Frigon *et al*., 2009; Hurteau *et al*., 2017; Mari *et al*., 2024). Here, we investigated the probability of evoking reflex responses in all four limbs before and after staggered hemisections by evaluating the distribution of SLRs (N1/P1), MLRs (N2/P2) and LLRs (N3/P3). We did this by calculating the fraction of the total number of SLRs or MLRs/LLRs separately, on recorded muscles for each limb across cats. For homolateral and diagonal responses in the left and right hindlimbs, we only evaluated MLRs/LLRs because of infrequent occurrence of SLRs. We excluded the H2T1 time point as only two cats were recorded.

We found no significant difference in response occurrence probability for left and right homonymous SLRs (left, p = .441, GLMM; right, p = .925, GLMM) and MLRs/LLRs (left, p = .496, GLMM; right, p = .401, GLMM) across states/time points (Fig. 6A). Similarly, the probability of evoking crossed SLRs (left, p = .085, GLMM; right, p = .304, GLMM) and MLRs/LLRs (left, p = .095, GLMM; right, p = .225, GLMM) in left and right forelimb muscles did not differ across states/time points (Fig. 6B). In contrast, we found a significant main effect of state/time point on homolateral MLRs/LLRs occurrence probability in the left (p = 1.00 × 10^-6^, GLMM) and right (p = 1.50 × 10^-5^, GLMM) hindlimbs (Fig. 6C). In the left hindlimb, homolateral responses were 8.3 (p = 7.00 × 10^-6^) and 22.7 (p = 1.25 × 10^-4^) times more likely to be evoked in the intact state compared to H1T1 and H2T2, respectively. They were also 5.2 (p = 3.73 × 10^-4^) and 14.4 (p = .001) times more likely to be evoked at H1T2 compared to H1T1 and H2T2, respectively. In the right hindlimb, homolateral responses were 10.0 (p = 1.00 × 10^-4^), 6.6 (p = 4.16 × 10^-4^) and 14.7 (p = 6.50 × 10^-5^) times more likely to be evoked in the intact state compared to H1T1, H1T2 and H2T2, respectively. We found a significant main effect of state/time point on diagonal MLRs/LLRs occurrence probability in the left (p = 7.12 × 10^-7^, GLMM) and right (p = 1.06 × 10^-9^, GLMM) hindlimbs (Fig. 6D). In the left hindlimb, diagonal responses were 18.2 (p = 8.00 × 10^-6^), 9.3 (p = 1.46 × 10^-4^) and 58.8 (p = 7.00 × 10^-6^) times more likely to be evoked in the intact state compared to H1T1, H1T2 and H2T2, respectively. They were also 6.3 (p = .036) times more likely to be evoked at H1T2 compared to H2T2. In the right hindlimb, diagonal responses were 28.6 (p = 4.84 × 10^-9^), 7.2 (p = 1.00 × 10^-5^) and 100.0 (p = 3.00 × 10^-5^) times more likely to be evoked in the intact state compared to H1T1, H1T2 and H2T2, respectively. They were also 4.0 (p = .009) and 13.8 (p = .015) times more likely to be evoked at H1T2 compared to H1T1 and H2T2, respectively.

**Figure 5.**
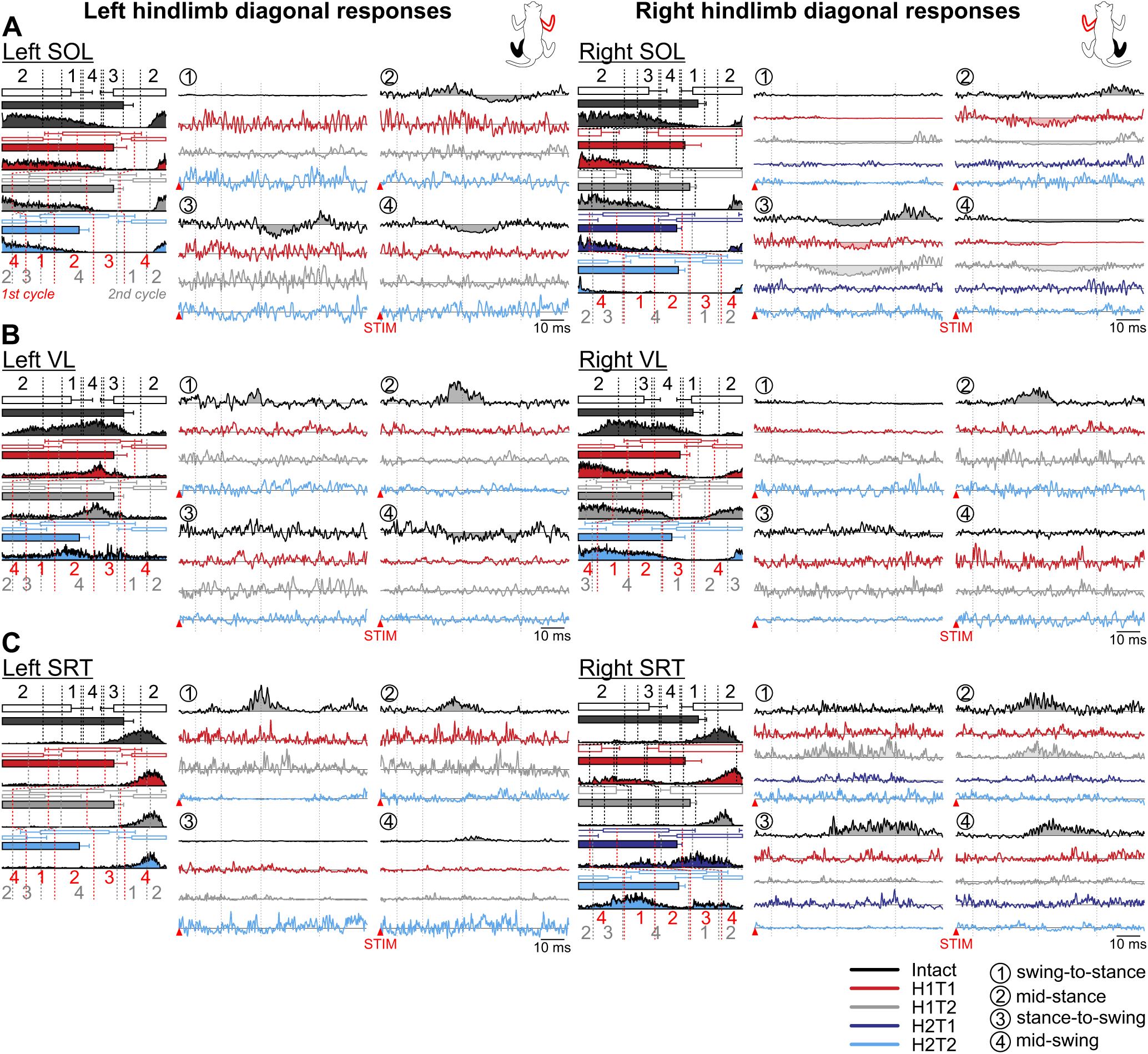
Phase-dependent modulation of cutaneous reflexes evoked in diagonal hindlimb muscles during locomotion before and following staggered hemisections. Each panel shows, from left to right, stance phases of the stimulated forelimb (empty horizontal bars) and diagonal hindlimb (filled horizontal bars) with its averaged rectified muscle activity normalized to cycle duration in the different states/time points, and diagonal reflex responses in representative cats for the left and right **(A)** soleus (LSOL, cat TO; RSOL, cat JA), **(B)** vastus lateralis (VL, cat TO), and **(C)** anterior sartorius (LSRT, cat TO; RSRT, cat JA). Reflex responses are shown with a post-stimulation window of 80 ms in four phases in the intact state, and after the first (H1) and second (H2) hemisections at time points 1 (T1) and/or 2 (T2). At each state/time point, evoked responses are scaled according to the largest response obtained in one of the four phases. The scale, however, differs between states/time points. The dotted vertical lines in the reflex responses indicate the N1/P1, N2/P2 and N3/P3 time windows.

**Figure 6.**
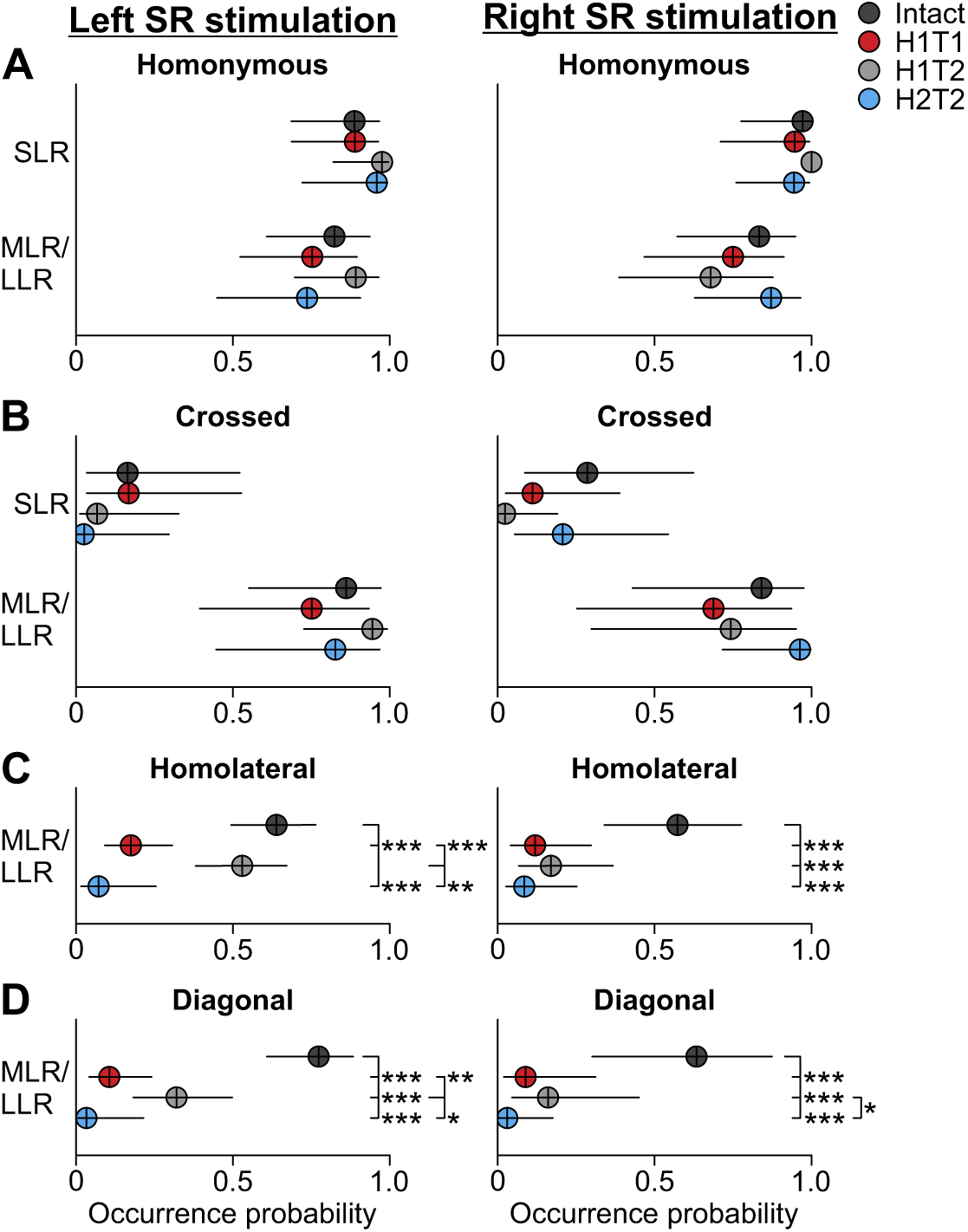
Reflex response occurrence in all four limbs before and after staggered hemisections. Response occurrence probabilities are shown for short- (SLR) and mid-/long-latency (MLRs/LLRs) responses with stimulation of the left or right superficial radial nerve before (intact) and after the first (H1) and second (H2) hemisections at time points 1 (T1) and/or 2 (T2). Tables 1, 2 and 3 provide details on the number of pooled data for SLRs, MLRs and LLRs. Each filled circle represents the mean probability ± confidence interval (95%) in 5 forelimb or 10 hindlimb muscles pooled across cats for homonymous/crossed (A and B) and homolateral/diagonal (C and D) responses, respectively. If a significant main effect of state/timepoint was found (generalized linear mixed model), we compared states/time points. Asterisks indicate significant differences at p < 0.05*, p < 0.01** and p < 0.001***. When one state/time point was significantly different from two states/time points, the comparison starts with a longer horizontal line.

Therefore, overall, after the first and second hemisections, the probability of evoking homolateral and diagonal MLRs/LLRs in hindlimb muscles with both left and right SR nerve stimulations was always lower compared to the intact state. In addition, with stimulation of the left SR nerve, the occurrence probability of evoking homolateral and diagonal MLRs/LLRS recovered after the first hemisection (at H1T2 compared to H1T1), before seeing a drastic decrease after the second hemisection.

## DISCUSSION

In the present study, we showed changes in reflex responses evoked by electrically stimulating cutaneous afferents of the forepaw dorsum (SR nerve stimulation) during locomotion after staggered hemisections, extending our recent study with reflex responses evoked by stimulating hindlimb cutaneous afferents in the same animals and lesion paradigm (Mari *et al*., 2024). The main result of the present study was a noticeable loss/reduction of mid- and long-latency homolateral and diagonal responses in hindlimb muscles, after both the first and second hemisections. However, after the first hemisection, we observed a partial recovery of these responses evoked by the left SR (contralesional) from the early to the late time point, which then disappeared after the second hemisection. These changes in homolateral and diagonal responses correlated with altered and weakened fore-hind coordination and impaired balance during quadrupedal locomotion, as we recently reported (Audet *et al*., 2023), and also discussed in relation with changes in reflex responses evoked by hindlimb cutaneous afferents (Mari *et al*., 2024). In the following sections, we discuss changes in reflex responses evoked by forelimb cutaneous afferents, the putative mechanisms and pathways involved and the functional significance of our results for locomotor control/recovery after SCI.

### Cutaneous reflexes from forelimb afferents before and after staggered lateral hemisections

In our staggered lateral hemisections paradigm, the first hemisection unilaterally disrupted direct descending motor pathways from the brain and cervical cord to the lumbar cord, as well as ascending pathways that carry somatosensory information from the hindlimbs to the cervical cord and then the brain. The second hemisection on the left side then disrupted these direct pathways contralateral to the first hemisection, generating a bilateral disruption. Lesion extent varied between animals (see Fig. 1B) and likely contributed to inter-individual variability. As expected, short- (N1/P1), mid- (N2/P2) and/or long-latency (N3/P3) responses in muscles of the homonymous and crossed forelimb remained after the first and second hemisections with both left and right SR nerve stimulations (Fig. 6). These responses also retained their phase-dependent modulation (**Table 1**). It is unlikely that the spinal lesions at T5-T6 and then T10-T11 damaged forelimb motoneuronal pools, which are located at C5-T2 spinal segments in cats (Sterling & Kuypers, 1967; Fritz *et al*., 1986*a*, 1986*b*; Hörner & Kümmel, 1993). The SR nerve originates from the brachial plexus, formed by the ventral branches of the last three cervical nerves and the first thoracic nerve, and enters spinal segments C7-T1 (Ansón *et al*., 2013; Mencalha *et al*., 2014; Hakkı Nur *et al*., 2020). Thus, the sensorimotor circuits responsible for homonymous and crossed reflex responses in forelimb muscles are largely preserved after thoracic SCIs.

Although the pattern of reflex responses in homonymous and crossed forelimb muscles remained generally similar after thoracic hemisections, we did observe several small changes (Figs. 2 and 3). These changes can involve supralesional circuit reorganization (Krupa *et al*., 2022) and/or the loss of inhibitory or excitatory ascending pathways from lumbosacral segments (Sterling & Kuypers, 1967; Giovanelli Barilari & Kuypers, 1969; Molenaar & Kuypers, 1978; English, 1985; Alstermark *et al*., 1987*b*; Dutton *et al*., 2006; Reed *et al*., 2006). Long ascending propriospinal neurons make direct contact with motoneurons in the most caudal cervical segments in rats (Reed *et al*., 2006, 2009; Brockett *et al*., 2013) and C3-C4 segments in cats (English, 1985; Alstermark *et al*., 1987*b*, 2007), with the vast majority being excitatory (Miller *et al*., 1973; Juvin *et al*., 2005; Reed *et al*., 2006; Brockett *et al*., 2013). These long projecting neurons play a role during locomotion as their silencing can disrupt left-right coordination of the hindlimbs and forelimbs (Pocratsky *et al*., 2020; Shepard *et al*., 2021). However, fore-hind coordination appears to mainly depend on short propriospinal neurons because blocking their activity at thoracic levels, without disrupting transmission in long propriospinal neurons, leads to independent rhythmic activity at cervical and lumbar levels in the in vitro neonatal rat preparation (Ballion *et al*., 2001; Juvin *et al*., 2005). Moreover, in both fictive and real locomotion studies in decerebrate cats with a complete thoracic transection performed at T10-T13, reflex responses evoked with SR stimulation were preserved and rhythmically modulated in forelimb motoneurons and muscles (Shimamura *et al*., 1990; Fuwa *et al*., 1991; Seki & Yamaguchi, 1997). Thus, the neural circuits modulating forelimb reflexes evoked by SR nerve afferents are mainly located at cervical and upper thoracic segments. This can include cervical spinal locomotor CPGs (Yamaguchi, 1992, 2004; Kinoshita & Yamaguchi, 2001) that interact with primary afferent inputs (Prochazka *et al*., 2002; Frigon & Rossignol, 2006; Frigon *et al*., 2021; Lalonde & Bui, 2021) and supraspinal signals (Shimamura & Livingston, 1963; Shimamura *et al*., 1984; Brink *et al*., 1985; Alstermark *et al*., 1987*a*; Fleshman *et al*., 1988; Fuwa *et al*., 1991; Bretzner & Drew, 2005; Ni *et al*., 2014; Bazley *et al*., 2014; Duysens, 2024). However, ascending pathways from the lumbosacral cord likely participate in part of the reflex response patterns and modulation, particularly in the intact state.

In hindlimb muscles, the occurrence of homolateral and diagonal mid- and/or long-latency reflex responses was considerably reduced after the first and second hemisections, with both left and right SR nerve stimulations (**Figs. 4-6**). These responses are thus highly dependent on the integrity of descending pathways running in the thoracic spinal cord. Thoracic lesions directly disrupt descending pathways from the brain and cervical cord that generate homolateral and diagonal reflex responses (Miller *et al*., 1977; Haridas & Zehr, 2003; Hurteau *et al*., 2018; Mari *et al*., 2023). Miller and colleagues (1977) proposed that long descending propriospinal neurons, with axons mainly traveling in the ventrolateral spinal cord (Flynn *et al*., 2011), constitute the main pathways responsible for homolateral and diagonal responses, although a contribution from short propriospinal neurons is also likely and cannot be excluded. Some of these pathways project ipsilaterally and/or contralaterally (Côté *et al*., 2018; Laliberte *et al*., 2019). Additionally, thoracic lesions disrupt brainstem pathways that release monoamines throughout the spinal cord and enhance neuronal excitability (Noga *et al*., 2009, 2011). Studies have shown that serotonin facilitates hindlimb cutaneous reflexes in spinal-transected cats (Barbeau & Rossignol, 1990) and rats (Maj *et al*., 1976; Nozaki *et al*., 1977).

We observed a partial return of the occurrence of homolateral and diagonal reflex responses evoked by the left SR nerve after the first hemisection, from the first to the second time point, consistent with ongoing neuroplasticity (Fig. 6). Thus, over the course of 6-7 weeks, changes occurred in spinal circuits to facilitate reflex transmission from cervical to lumbar levels. Numerous studies have shown that the spinal neural circuitry below an incomplete SCI undergoes substantial reorganization (Barriere *et al*., 2008; Frigon *et al*., 2009; Barrière *et al*., 2010; Martinez *et al*., 2011, 2012; Gossard *et al*., 2015), including after staggered lateral hemisections (Jane *et al*., 1964; Kato *et al*., 1984, 1985; Stelzner & Cullen, 1991; Courtine *et al*., 2008; Van Den Brand *et al*., 2012; Cowley *et al*., 2015; Audet *et al*., 2023). After the first hemisection, the left side of the spinal cord remains relatively intact, and plasticity can occur in descending pathways, which can be beneficial or detrimental (Beauparlant *et al*., 2013; Fink & Cafferty, 2016; Shepard *et al*., 2023). This can involve establishing new connections and/or reactivating latent ones, as well as reinforcing spared descending propriospinal and supraspinal pathways through spontaneous collateral sprouting that target both lesioned and unlesioned fibers (Edgerton *et al*., 2004; Cai *et al*., 2006; Maier & Schwab, 2006; Fenrich & Rose, 2009; Basaldella *et al*., 2015; Higgin *et al*., 2020; Zavvarian *et al*., 2020). The return of neuronal excitability at lumbar levels could have also facilitated responses evoked by descending pathways.

The recovery of homolateral and diagonal responses evoked by stimulating the left SR from the early to the late time point after the first hemisection was lost after the second hemisection, which disrupted descending pathways on the left side. This staggered thoracic lateral hemisections paradigm reveals limitations to the formation of new short connections to enable reflex transmission from cervical to lumbar levels. Previous studies have shown that new propriospinal relays can form spontaneously to reroute the supraspinal influences onto lumbar circuits (Bareyre *et al*., 2004; Courtine *et al*., 2008; Cowley *et al*., 2008, 2010; Zaporozhets *et al*., 2011; May *et al*., 2017). However, these new connections appear limited in supporting cutaneous reflex transmission from cervical to lumbar segments. This is consistent with studies using the in vitro neonatal rat brain stem-spinal cord preparation that required neurochemical excitation of thoracic propriospinal neurons to generate hindlimb locomotion with brainstem electrical stimulation after staggered thoracic lateral hemisection (Cowley *et al*., 2008; Zaporozhets *et al*., 2011). Figure 7 schematically presents a scenario explaining changes in reflex responses to the four limbs after staggered hemisections with left and right SR nerve stimulation.

**Figure 7.**
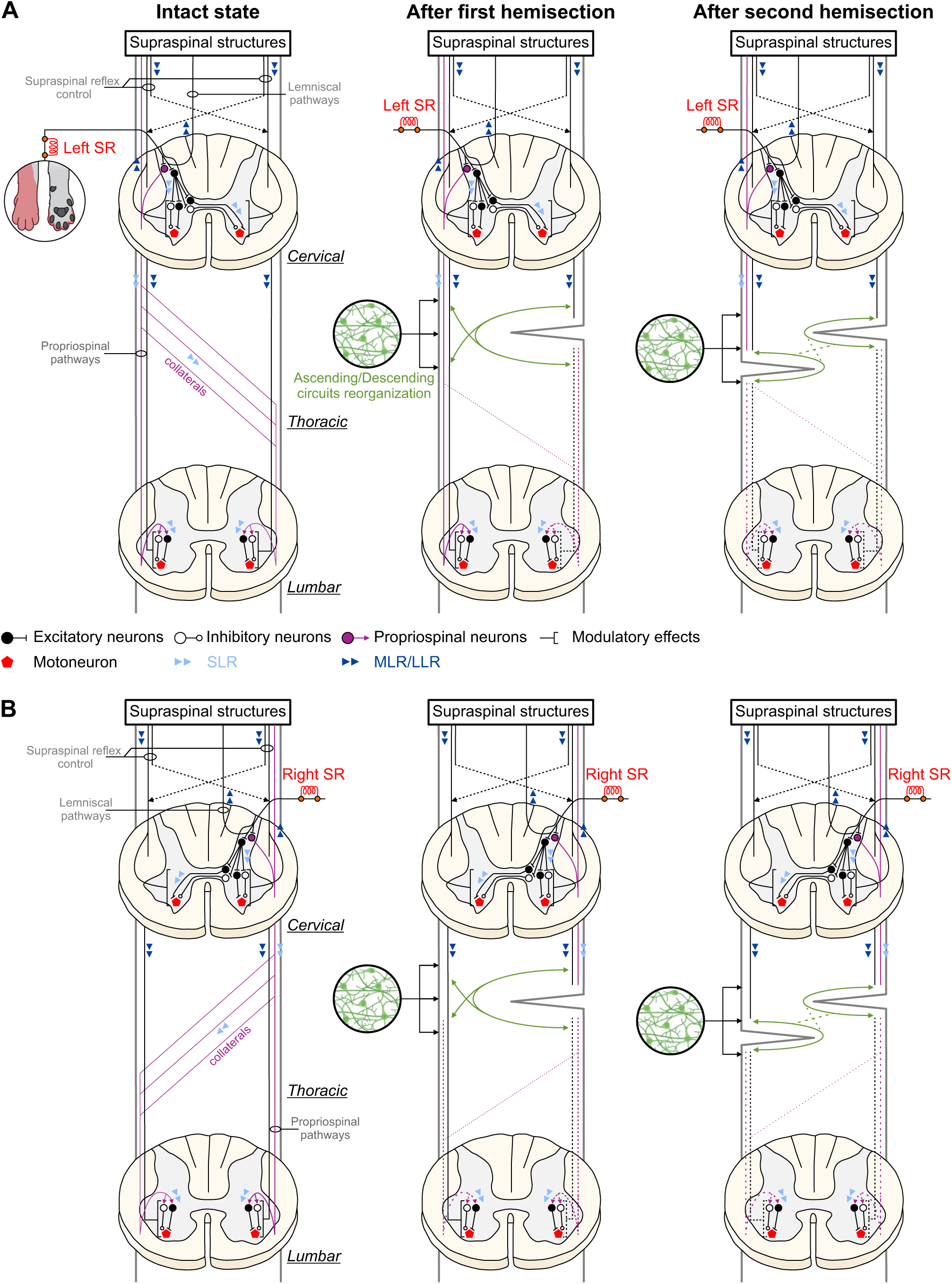
Schematic illustration of putative pathways and mechanisms contributing to cutaneous reflexes and their modulation before and after staggered hemisections. In the intact state, afferents from the left **(A)** and right **(B)** superficial radial (SR) nerves contact spinal interneurons that project to motoneurons within the hemisegment (homonymous responses) at cervical levels and commissural interneurons projecting contralaterally (crossed responses). SR nerve afferents also make contacts with propriospinal neurons that project to lumbar levels, terminating ipsilaterally (homolateral responses) or contralaterally via collateral projections at different segments (diagonal responses). The pathways responsible for short-latency responses (SLRs) are mainly confined to spinal circuits, including SLRs in hindlimb muscles. The pathways contributing to mid- and long-latency responses (MLRs/LLRs) transmit sensory information to supraspinal structures via long ascending projection neurons (propriospinal and/or dorsal lemniscal pathways) that project back to spinal circuits controlling the fore- and hindlimbs. After the first hemisection (on the right side), SLRs and MLRs/LLRs in forelimb muscles remain present although their response pattern can change due to the loss of inhibitory or excitatory ascending pathways from lumbosacral circuits. The occurrence of MLRs/LLRs in hindlimb decreases (dashed lines) due to disruptions in ascending and descending pathways to and from supraspinal structures. Spared supraspinal axons are potentially strengthened or sprout to form new connections to transmit descending signals. After the second hemisection (on the left side), direct ascending and descending pathways are both disrupted, and reorganization of short propriospinal neurons is required to relay signals through the lesions, although their ability to do so is limited, leading to a considerable loss in MLRs and LLRs in hindlimb muscles.

### Functional considerations

Electrically stimulating the SR nerve mimics a mechanical contact of the forepaw dorsum, eliciting a functional response consistent with a stumbling corrective or preventive reaction during the forelimb swing and stance phases, respectively, as shown in intact and decerebrate cats (Miller *et al*., 1977; Matsukawa *et al*., 1982; Drew & Rossignol, 1985, 1987; Shimamura *et al*., 1990; Fuwa *et al*., 1991; Hurteau *et al*., 2018; Mari *et al*., 2023). In the present study and other studies, stimulating the SR nerve evoked reflex responses in muscles of the four limbs in intact cats, consistent with a whole body functional response to an external perturbation to alter limb trajectory and ensure dynamic balance (Haridas & Zehr, 2003; Hurteau *et al*., 2018; Pearcey & Zehr, 2019*a*; Merlet *et al*., 2022; Mari *et al*., 2023). Our results indicate that functional responses to SR nerve stimulation would have been appropriate in forelimb muscles to alter the trajectory of the ipsilateral forelimb and reinforce support in the contralateral forelimb after staggered thoracic hemisections, as reflex responses in forelimb muscles and their phase modulation were largely preserved. However, the loss/reduction of homolateral and diagonal responses suggest an inability to coordinate the hindlimbs, which likely would have affected the whole-body response to a real perturbation.

The loss/reduction of homolateral and diagonal responses also reflects a disruption of neural communication between the brain/cervical cord and the lumbar cord, which cannot be compensated by new connections, as discussed above. This affects sensorimotor functions that depend on this communication. Indeed, after staggered thoracic hemisections, we recently showed that fore-hind coordination was altered and weakened (Audet et al. 2023). Moreover, after the second hemisection, balance assistance was required during treadmill locomotion. Although we can only speculate, the loss of cutaneous reflex transmission from cervical to lumbar levels could have contributed to weakened interlimb coordination and impaired postural control.

In humans, cutaneous afferents from the arm contribute to locomotor and/or postural control. For instance, cutaneous stimulation at the wrist at the swing-to-stance transition during walking results in increased ankle dorsiflexion (Haridas & Zehr, 2003). This could slow forward progression by reducing propulsion from ankle plantarflexors, thereby minimizing a possible collision with an object with the outstretched arm. Leg responses evoked from hand cutaneous nerves were facilitated with vision removed during walking, but were restored when participants were asked to lightly touch a stable reference (Forero & Misiaszek, 2015) that reinforced stability (Dickstein & Laufer, 2004; Forero & Misiaszek, 2013). Thus, forelimb cutaneous afferents can assist with balance during walking. In cats, more weight is distributed to the forelimbs for stability and propulsion after spinal lesions (Rossignol *et al*., 1999) and maintaining proper sensorimotor interactions in cervical sensorimotor circuits is important, as shown in the present study.

Another factor to consider after single or staggered thoracic lateral hemisections is the emergence of spatiotemporal left-right asymmetries between the hindlimbs (Kato et al., 1985; Martinez et al., 2011, 2013; Audet et al., 2023), which can lead to walking instability (Dambreville *et al*., 2015; Huijben *et al*., 2018). Recent studies using split-belt locomotion have shown that inducing left-right asymmetries reduced hindlimb reflex responses in some muscles with SR or SP nerve stimulation (Hurteau *et al*., 2017; Hurteau & Frigon, 2018; Mari *et al*., 2023). Split-belt locomotion induces left-right asymmetries in sensory feedback from the limbs, with increased loading for the limbs on the slow belt (Frigon *et al*., 2015; Park *et al*., 2019). The hemisections also induce left-right asymmetries as the contralesional hindlimb spends a greater proportion of the cycle supporting bodyweight (Martinez *et al*., 2011, 2012; Audet *et al*., 2023). This increased loading generates asymmetric limb sensory feedback, which could play a role in modulating interlimb reflexes.

### Methodological limitations

A limitation of the present study was that balance assistance was required after the second hemisection to conduct reflex testing during locomotion, with an experimenter holding the tail for medio-lateral stability, but without providing weight support, as discussed in our recent study (Mari *et al*., 2024). Without this balance assistance, cats stumbled every few steps. Balance assistance likely facilitated reflex responses and their phase-dependent modulation but without it, we could not have conducted reflex testing and compared it to the other states (i.e. intact and following the first hemisection). Another limitation is that we pooled reflex responses in hindlimb muscles according to the phase of the stimulated forelimb. As stated, cats often performed 2:1 fore-hind patterns after the first and second hemisections, where a forelimb performed two cycles within a hindlimb cycle. Because our stimulation is based on the forelimb cycle, it means that after spinal lesions, some stimuli occurred in different phases of the hindlimb cycle. We acknowledge this as a limitation but separating all responses so that both forelimb and hindlimb phases corresponded would have required many more stimuli and several more minutes to complete a session. These studies are difficult to perform in spinal cord-injured cats. Thus, for a given forelimb phase of stimulation, we pooled all responses evoked in a hindlimb muscle, irrespective of the phase of the hindlimb. This could have affected average responses, particularly if inhibitory and excitatory responses occurred at the same latency in different hindlimb phases after hemisections. However, we investigated individual responses and observed that most homolateral and diagonal responses were simply lost after hemisections.

### Concluding remarks and clinical perspectives

In conclusion, the main finding of this study was the loss/reduction of homolateral and diagonal reflex responses in hindlimb muscles from forelimb cutaneous afferents after staggered thoracic hemisections, which correlated with weakened coordination between the fore- and hindlimbs and impaired balance. How do our findings in the cat model relate to people with spinal cord injury? Our paradigm offers a substrate to test the efficacy of therapeutic approaches (e.g. spinal cord stimulation, pharmacology) to restore neural communication between the spinal locomotor networks controlling the arms/forelimbs and legs/hindlimbs. Exploiting cervicolumbar connections, with rhythmic arm movements and/or primary afferent stimulation, could facilitate locomotor rehabilitation after SCI. For instance, in cats (Harnie *et al*., 2024) and humans (Frigon *et al*., 2004; Zehr *et al*., 2004; Hiraoka & Iwata, 2006; Loadman & Zehr, 2007; Javan & Zehr, 2008; Dragert & Zehr, 2009; Hundza & Zehr, 2009; De Ruiter *et al*., 2010; Hundza *et al*., 2012; Massaad *et al*., 2014; Pearcey & Zehr, 2019*b*), rhythmic arm movements contribute and/or modulate reflexes in leg/hindlimb muscles. In rats, quadrupedal locomotor training improved the quality of fore-hind coordination after a thoracic spinal cord hemisection, with recovery correlating with an increased number of propriospinal neurons just above and below the injury site (Shah *et al*., 2013). In humans with incomplete SCI, coordinating rhythmic arm movements simultaneously with the legs facilitates corticospinal and/or corticofugal drive (Zhou *et al*., 2017) to leg muscles and modulates cervicolumbar connectivity (Zhou *et al*., 2018). Combined arm and leg movements improved locomotor function. Applying transcutaneous spinal cord stimulation at the cervical levels can also modulate the activity of lumbar networks (Barss *et al*., 2020; Parhizi *et al*., 2021). Therefore, these results highlight the importance of activating forelimb sensory feedback, engaging cervical locomotor networks and descending propriospinal pathways in the recovery of leg/hindlimb locomotor movements after SCI.

## Support or grant information

This work was supported by a grant from the National Institutes of Health: R01 NS110550 to AF, IAR and BIP. AF is a Fonds de Recherche-Santé Quebec (FRQS) Senior Research Scholar. JA and JH were supported by FRQS doctoral scholarships and ANM by a FRQS postdoctoral scholarship.

## Author contributions

SM, IAR, BIP, and AF contributed to conception and design of the study. SM, CL, AM, JA, SY, RA and JH conducted the research. SM organized the database and performed the data and statistical analysis. SM and AF wrote the first draft of the manuscript. All authors contributed to manuscript revision, read, and approved the final version.

## Acknowledgments

We thank Philippe Drapeau for providing data acquisition and analysis software, developed in the Rossignol and Drew laboratories at the Université de Montréal. We thank the Biostatistics department of the Centre de Recherche du Centre Hospitalier Universitaire de Sherbrooke for statistical assistance.

## Data availability statement

The raw data supporting the conclusions of this article will be made available by the authors, without undue reservation.

## Ethics statement

The animal study was reviewed and approved by the Animal Care Committee of the Université de Sherbrooke.

## Competing statement

The authors declare no competing financial interests.

